# When anticipation is not enough: a mixture of robust and adaptive feedback control strategies improve reaching in dynamic environments

**DOI:** 10.64898/2026.04.07.716902

**Authors:** Hari Teja Kalidindi, Frédéric Crevecoeur

**Affiliations:** Institute of Communication Technologies, Engineering and Applied Mathematics, UC Louvain, Louvain-la-Neuve, Belgium; Institute of Neuroscience, UC Louvain, Brussels, Belgium; WEL Research Institute, Wavre, Belgium; Donders Center for Cognition, Radboud University, The Netherlands

## Abstract

Successful goal-directed movements depend on the central nervous system’s (CNS) ability to handle diverse physical interactions. The CNS is thought to handle different dynamical contexts through three mechanisms: (i) trial-by-trial adaptation when forces are predictable, (ii) a model-free robust control strategy, and (iii) online adaptation of feedback responses. While each has been studied independently, their relative contributions and the possibility that they are recruited to different extents across contexts is unknown. Here, we quantified all three strategies within the same individuals to examine how CNS exploits them under varying environmental conditions. Participants (19 female, 15 male) performed reaching tasks while interacting with robot-generated force-fields that were either consistent or varied unpredictably. Trial-by-trial adaptation was measured using standard force channels to isolate anticipatory compensation. Robust control was assessed through movement velocity and corrective force magnitude. Online adaptive control was quantified by the temporal alignment between commanded and measured forces within a movement. Results showed that participants improved anticipatory compensation in consistent environments and relied on both robust and online adaptation when perturbations were unpredictable. Crucially, markers of robust control dominated the early movement phase, whereas online adaptation dominated later corrections. This temporal dissociation was confirmed by electromyographic recordings. Markers of robust and online adaptive feedback strategies also statistically predicted participants’ ability to adapt across trials in consistent environments, revealing a common trait linking online control and adaptation. These findings reveal a rich and flexible combination of control mechanisms, offering a new framework for understanding the neurophysiological bases of reaching control.

**Significance Statement:** Human reaching control is a complex behavior resulting from several mechanisms that orchestrate feedback responses to mechanical perturbations and adaptation to changes in the environment. Here we combine previously studied paradigms to highlight within the same groups of healthy volunteers that three major components are recruited to different extents dependent on the context: unpredictable environment promote concomitant use of robust control and online adaptation whereas predictable environments recruit standard adaptation based on anticipatory compensation. Remarkably, individuals’ adaptive capabilities correlated across consistent and inconsistent environments, suggesting a key involvement of adaptive mechanisms in both online control and trial-by-trial adaptation. Robust control, online adaptation, and anticipatory compensation are dissociable behaviorally, and are used to varying levels as a result of individual traits.

## Introduction

Humans exhibit a remarkable ability to interact with their environment. This proficiency is often attributed to our ability to form internal representations of movement dynamics, known as internal models, which allow us to anticipate interactions and plan motor commands (Franklin and Wolpert, 2011). This anticipatory strategy commonly described as feedforward control, is effective when changes in dynamics are consistent and predictable across movements (Gonzalez Castro et al., 2014). However, when faced with novel or variable environmental forces – for instance, the swaying of a moving bus – errors in internal models may induce movement deviations from the intended goal, requiring adjustments also in feedback control. A key open question is how the CNS maintains performance in such unpredictable scenarios (Kalidindi and Crevecoeur, 2023).

Recent research has identified two distinct strategies for improving feedback control (Figure 1). The first is a model-free robust control strategy (Crevecoeur et al., 2019; Bian et al., 2020), in which the CNS does not rely on a precise model of potential environmental interactions. Instead, it increases control gains with uncertainty of disturbances. In the context of human reaching, this strategy leads to stronger corrective responses to counteract any unmodelled disturbances (Crevecoeur et al., 2019). Motor control in variable environments has been linked to increased grip force (Hadjiosif and Smith, 2015), upregulated visuomotor and proprioceptive feedback gains (Franklin et al., 2012, 2017; Maurus et al., 2023), heightened muscle stretch responses (Crevecoeur et al., 2019; Coltman and Gribble, 2020), and faster movements (Izawa et al., 2008; Crevecoeur et al., 2019). These observations may all reflect a robust control strategy as they involve an upregulation of feedback gains impacting both goal-directed motor commands and feedback responses to disturbances.

**FIGURE 1.**
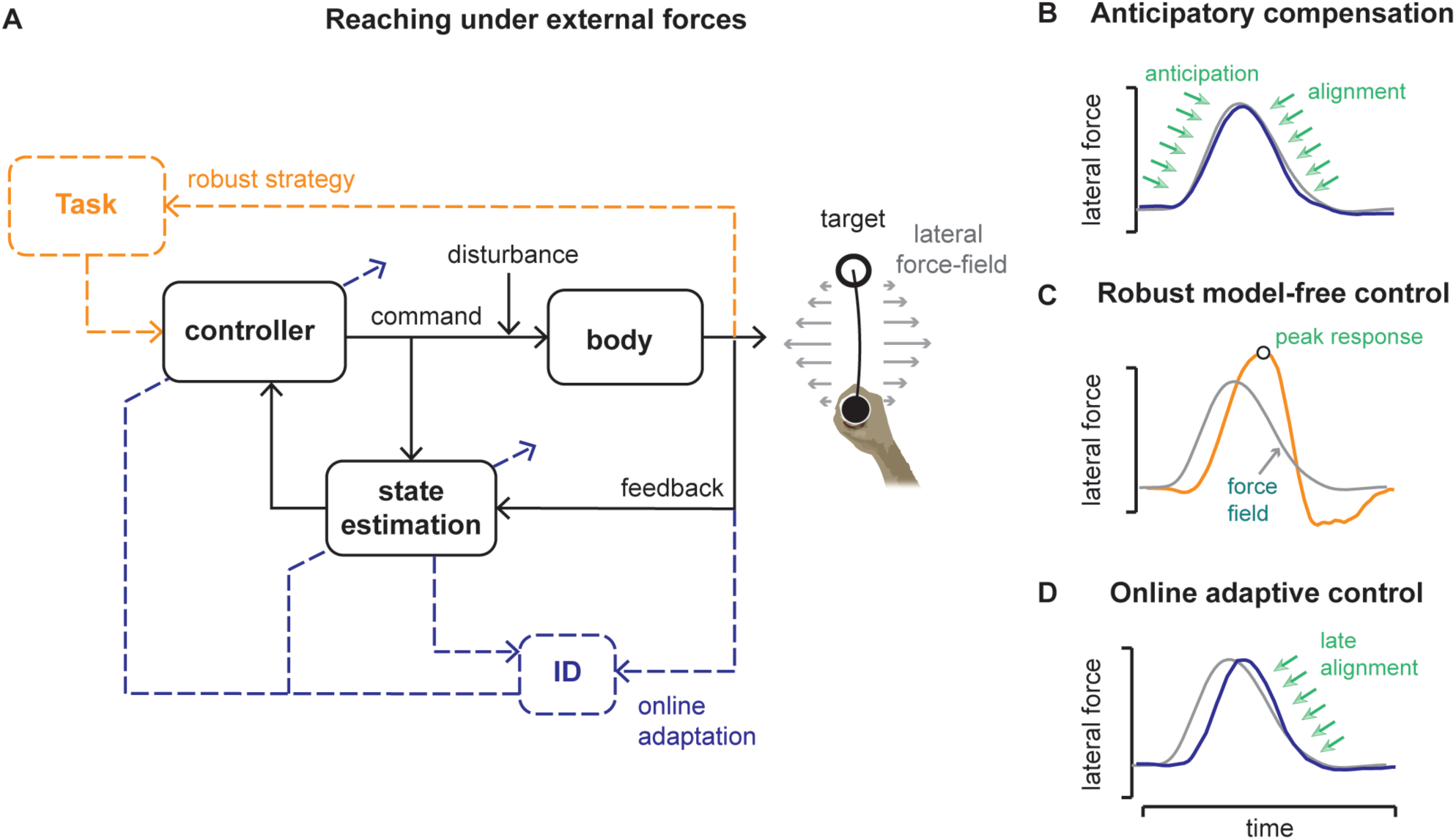
Different control modes interact to produce reaching in stochastic environments. A) for reaching in variable environments two distinct types of adjustments in feedback control have been reported, that includes robust and adaptive feedback responses B) when perturbations are predictable, humans can anticipate the forces and apply appropriate early corrections C) under novel, unpredictable environments robust responses involve increase in feedback gains and hence higher corrective forces D) adaptive feedback responses improve the alignment between hand applied forces and perturbation in the late movement period. The ID box illustrates the identification of model parameters used to update the controller. The nominal optimal feedback control architecture is represented in black; the system identification (ID) or updates in the internal model parameters and its influence on the controller and state estimation is in dashed blue lines; the updates in controller with a robust strategy is in dashed orange lines.

The second strategy involves online adaptation: the internal model is updated during movement based on the errors between predicted and actual sensory feedback processed in real-time (Braun et al., 2009; Crevecoeur et al., 2020b, 2020a, 2022; Mathew et al., 2020; Mathew and Crevecoeur, 2021; Doyen et al., 2025). The updated model allows corrective forces to align with encountered perturbations as the movement unfolds. Evidence from reaching tasks with unpredictable force-field perturbations shows that the corrective forces align with the perturbation later within the trial (Crevecoeur et al., 2020b, 2020a). Notably, online adaptive control is clearly dissociable from “nominal” feedback control – where the controller is based on fixed set of parameters (Todorov and Jordan, 2002; Todorov, 2005) – as online modification of the internal model alters the shape of the online feedback correction.

These strategies reflect a fundamental trade-off: robust control favors stability at the cost of increased motor output, whereas adaptive control promotes efficiency at the cost of sensitivity to internal model errors. Robust and adaptive control are both in principle useful to counter unmodelled disturbances, in the sense that movement deviations result from errors in the internal model. Besides these strategies, it was also suggested that limb mechanical impedance is increased to counteract perturbations at the muscle-periphery level (Burdet et al., 2001; Franklin et al., 2008). However, empirical measures of limb impedance maybe overestimated and overlap with the influence of CNS-mediated feedback responses (Crevecoeur and Scott, 2014). Overall, it remains unclear how the CNS selects or combines these different feedback control strategies depending on the context.

To investigate this, participants performed horizontal reaching movements towards visual targets while interacting with robotic-generated force-fields. The direction of the force-field was randomized across trials, creating environments with varying predictability. We examined how participants modulated control by analyzing kinematics, hand forces, and surface electromyography (EMG). Specifically, we aimed to determine whether feedback gains (used as proxy of robust control) and within-trial force alignments (indicative of adaptive control) were dissociable, and how they influenced the overall movement performance. Our results show that these behavioural markers were indeed dissociable, and our analyses further indicated precisely timed contributions of these feedback responses to counteract unpredictable forces.

## Materials and Methods

### Experiment Design

Thirty-four healthy participants (19 females) were recruited for the two experiments. All participants were right-handed and had no known history of any neurological disorder. Participants gave informed consent and the experiments were approved by the ethics committee at the host institution (UCLouvain, Belgium).

Movements consisted of 15 cm forward reaches toward visual targets (Figure 2). Both experiments were performed with a KINARM robotic manipulandum (KINARM, Ontario, Canada), and the parameters of the protocol described here were common across the different conditions and experiments. Participants were instructed to grab a robotic handle and move a hand-aligned cursor to the home target (radius 0.6 cm) projected onto a horizontal screen. The home target was initially red and turned green when the cursor entered the target. The goal target was presented as an open circle (radius: 1.2 cm) with a white edge, located 15 cm directly in front of the home target along the y-axis (see Figure 2A, B). After a random delay following stabilization in the home target (between 2 and 4 s, uniformly distributed), the goal circle was filled in with red color, coincident with a 100ms beep sound from surrounding speakers, providing a “go signal” for participants to begin their movement. If participants reached the goal target in less than 0.6 s following the go cue (including reaction time), the target turned back to an open circle to indicate that they reached it too soon. If they reached the goal target after 0.8 s, the target remained red indicating that they took too long. The goal target turned green when participants reached the goal target between 0.6-0.8s. The trial was considered successful if the hand-aligned cursor remained stable in this target for 1 s. There was no constraint on the movement speed, only on the arrival time, including reaction time, was constrained. These instructions were used to encourage similar movement timing and speed, but all trials were included in the dataset. The variations in protocols for the two experiments is described below.

**FIGURE 2.**
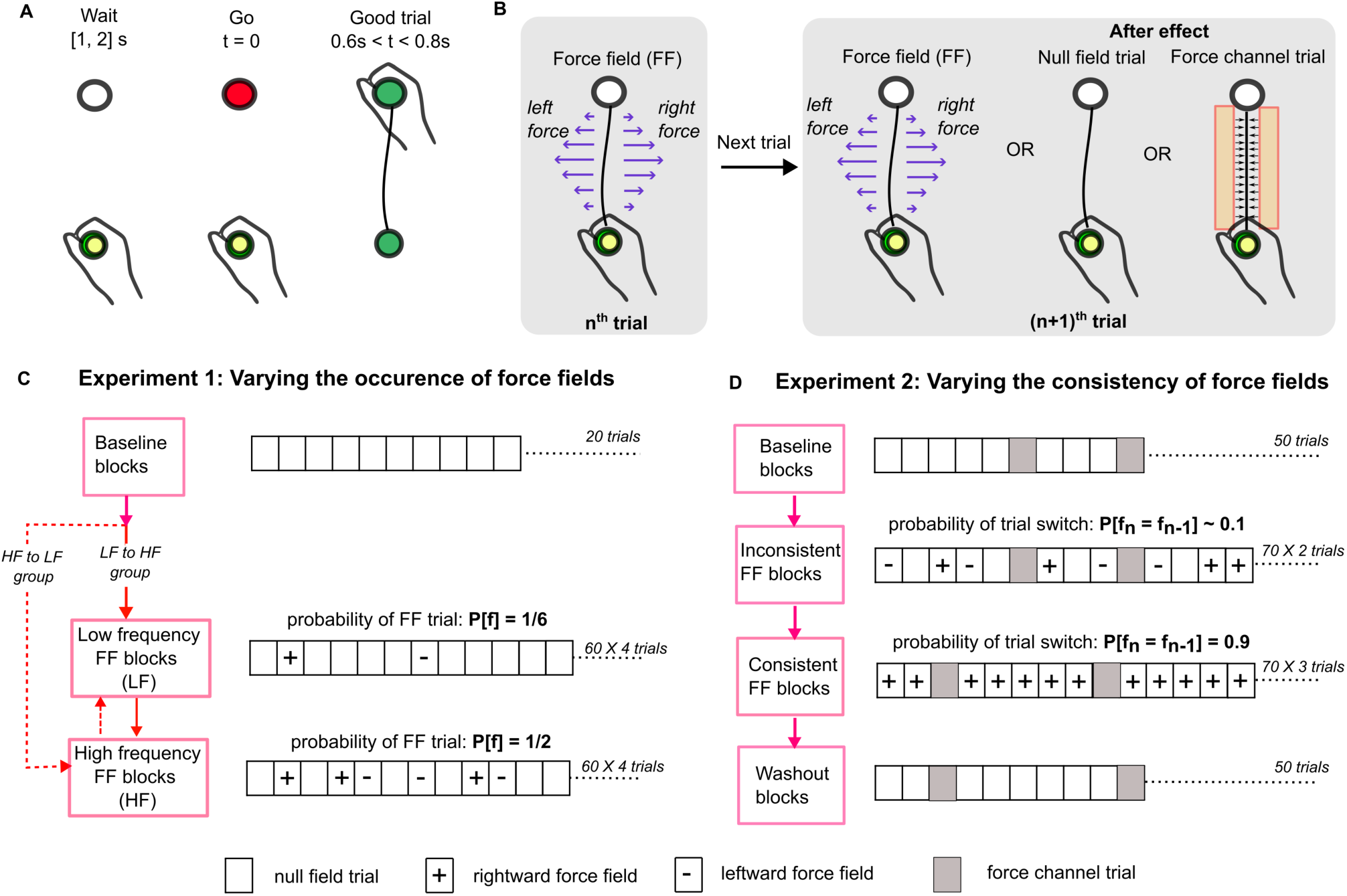
Schematic illustration of experiments. A) typical trial B) switching force field dynamics from one trial to the other, where a given null, force channel, rightward, and leftward force field (FF) trial can be followed by any possible trials. C) different conditions within Experiment 1, where participants were evenly divided to either start with LF or HF conditions D) different conditions within Experiment 2, where force channels were used to measure the anticipatory responses following standard procedures.

#### Experiment 1

The purpose of this experiment was to quantify, within the same participants, the impact of unpredictable changes in environment dynamics on their control strategies. Participants (n=17) performed a practice series of 20-30 trials in the null field, without any perturbation, until they felt comfortable with the task and instructions. Then, they performed 50 null-field trials referred to as the baseline condition. Participants were explicitly told they would not encounter any perturbation during the movement. These trials corresponded to pre-exposure phase performed before the introduction of trials with mechanical disturbances. This phase was followed by two different conditions presented sequentially, differed in the frequency of force field trials applied randomly amid null field trials. We refer to these conditions as High Frequency (HF) or Low Frequency (LF). Participants were randomly assigned to one of two groups that performed either the LF condition first, followed by the HF condition, or vice-versa (see Figure 2C).

Each of the HF and LF condition comprised 4 blocks of 60 trials. In LF condition, 60 trials in each block were divided into 50 null field trials and 10 force field trials (probability of a force field in a trial, p(f) = 1/6), which consisted of orthogonal force fields randomly interleaved (5 rightward, and 5 leftward). The orthogonal force field was defined as a lateral force along the *x*-axis proportional to the forward hand velocity as follows;

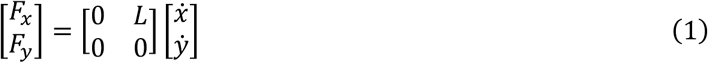

where the *x* and *y* coordinates correspond to lateral and forward axes relative to the reach path, *F_x,y_* are the force components along each axis and the dots indicate time derivatives. The value of *L* was ±13 Ns/m for rightward or leftward perturbations, respectively.

In HF condition, the 60 trials in each block were composed of 30 null field trials, 10 trials of orthogonal force fields (5 per direction) as described above, 10 trials of curl force fields (5 per direction), and 10 trials with lateral step perturbation loads (5 per direction). Overall, in the HF condition, the probability of a perturbation was *p*(*f*) = 0.5.

The curl fields were defined as follows:

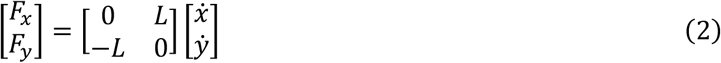

where *L* = ±13 *Ns*/*m*. The step perturbation had a rise time of 10ms and a magnitude of ±9 N. Notably, the orthogonal force field trials in HF conditions were the same as in the LF condition.

#### Experiment 2

Experiment 1 focused on the modulation of robust and adaptive control strategies across conditions of uncertainty, without any possibility to use trial-by-trial adaptation due to the unpredictable nature of the conditions. Here we designed the second experiment specifically to dissociate these components from the standard trial-by-trial anticipatory compensation observed in consistent environment (Figure 2D). This experiment was also designed to measure the transition between environments in terms of movement strategies. A different group of 17 healthy participants (9 women) were enrolled in it. On a given trial, during the reaching movement the manipulandum either applied no force (null field trial), a rightward or a leftward orthogonal force field, or a force channel used to measure anticipatory compensation for the force field based on standard techniques (Figure 2B) (Smith et al., 2006). During these channel trials, the robot motors were used to constrain movements to a straight path between the start position and the goal target. This channel was ∼1 mm wide, with a virtual wall with stiffness parameter -6,000 N/m and a damping coefficient of -50 Ns/m. Force channel trials acted as error-clamp trials where movement errors were removed by preventing any cursor motion perpendicular to the target direction (lateral *x* direction). As any lateral deviations and their online corrections are eliminated, it is commonly assumed that the force output on the lateral walls from these trials reflects the participants’ anticipatory compensation for the externally applied force field learnt from previous trials.

This experiment comprised 450 trials divided into 4 different conditions. It began with a baseline condition where participants made 48 trials in the absence of force fields (null fields), and 2 force channel trials interleaved in the last 10 null field trials in this condition (Figure 2D). This was followed by an inconsistent force field (random) condition (2 blocks with 70 trials per block). The 140 trials of this condition were composed of 40 null field, 40 rightward and 40 leftward force field trials, plus an additional 20 force channel trials. All trial types were randomly interspersed. The inconsistent condition was then followed by a consistent force field condition (3 blocks of 70 trials per block, totaling 210 trials). The total 210 trials in this condition comprised 180 force field trials applied only in the rightward direction, and 30 randomly interleaved force channel trials. Finally, a washout condition composed of 50 trials of null field trials and 10 randomly interleaved force channel trials were used in the end of the experiment.

Importantly, both the random and consistent conditions were constrained to have different lag-1 autocorrelation structure, the chance that a given trial type be experienced again in the next trial. This is enforced by a single parameter, which is the probability that the current trial is of the same type as the previous trial. In the inconsistent condition, this probability was 0.1. In the consistent condition, this probability was 0.9:

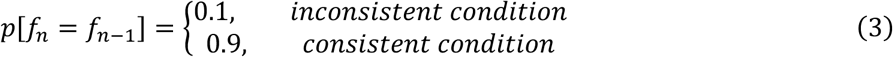

Note that in the consistent condition, the probability of trial type consistency was slightly less than 1 because of the presence of force channel trials interleaved between force field trials. To generate such a consistent environment, we first generated a randomly interleaved array of force field and channel trials, and then iteratively shuffled by swapping random pairs of differing trial types until the probability estimated from Equation 3 converged to 0.9.

The order of force field and force channel trials was pseudorandomized such that each trial type could not be predicted, and every participant experienced the trials in the same predetermined order. In the consistent condition the participants were not explicitly told that the force field would occur in the same direction such that their behaviour resulted from the sensorimotor experience of this block. In the baseline and washout conditions, the participants were explicitly told that no lateral perturbation would be applied.

### Data collection and Analysis

The two-dimensional coordinates of the hand-aligned cursor and components of the endpoint force were sampled at 1 kHz. The cursor velocity was obtained numerically with a fourth-order central-differences algorithm. In both experiments, we recorded the activity of mono-articular shoulder muscles (PM: pectorialis major and PD: posterior deltoid), known to be strongly recruited by lateral disturbances with the same setup and perturbation types (Crevecoeur et al., 2019, 2020a; Kalidindi and Crevecoeur, 2024). The electrodes were attached to the skin above the muscle belly after light abrasion with alcohol. The signal was amplified (gain: 104), digitally bandpass filtered with a dual-pass, fourth-order Butterworth filter (10-450 Hz), and normalized to the average activity across 0.5 s recorded when participants maintained postural control at the home target against a background load of 12N applied three times in four directions (forward, backward, right, left). This calibration block was performed at the end of first and eighth blocks in Experiment 1 and at the end of first and sixth (final) blocks in Experiment 2. We extracted the following parameters on a trial-by-trial basis:

#### Hand path error

Hand path error is used to quantify the total lateral deviation (*x*) of the hand path from a straight line connecting the start and the target locations for a given duration (*T*). Essentially, path error computes the root mean square hand deviation over the reach duration and is expressed in centimeters (*cm*).

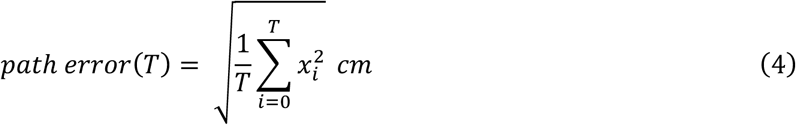

Notably, path error has been computed separately for rightward and leftward force field trials in Experiment 1, and the average between these two force field directions has been depicted in Figure 3A.

**FIGURE 3.**
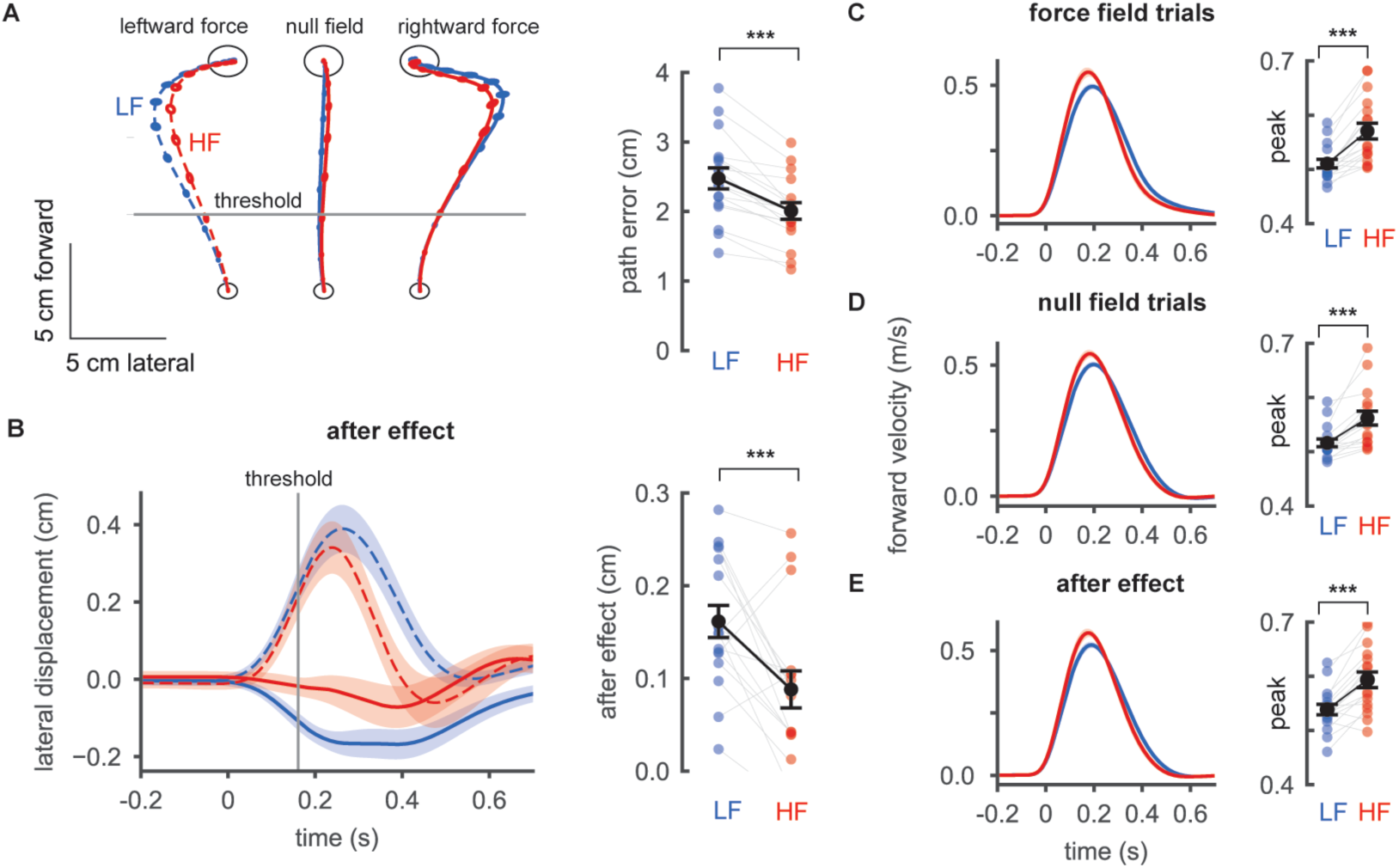
Kinematics of reaching movements in different contexts. A) group average hand path in the unperturbed trials within high frequency (HF, red) versus low frequency (LF, blue) perturbation scenario. The path error quantifies the root mean square lateral deviation of the hand path from a straight line connecting the start and targets. Dashed lines are hand path displacements during leftward force trials B) group average of hand path in the null field trials that immediately followed a force field trial. The after effect quantifies the lateral hand deviation at the threshold (see Methods). Dashed lines are hand path displacements in the null-field trial following a leftward force-field C) average forward hand velocity in the orthogonal force field trials in LF and HF conditions. D) same as (C) across null field trials E) same as (C) for the null field trials that immediately follow force field trials. Each dot in the error bar plot represents individual participant and gray lines reflect change in individual data across conditions. Vertical bars represent the S.E.M across participants (n=17). ***p<0.001

#### Measure of robust responses

It was earlier shown in (Crevecoeur et al., 2019) that the robust feedback responses in reaching control could be linked to an increase in peak hand velocity in the unperturbed directions, and an increase in lateral force applied by the participants to counteract disturbances. In Experiment 1, we used peak lateral force in the trials with force fields, and peak forward velocity as a proxy of change in control strength assuming that they were indicating the robustness of the controller. In Experiment 2, we quantified changes in control strength in terms of peak lateral force, while similar trends were observed for peak forward velocity.

#### Measure of online movement adaptation (Alignment index)

To quantify the online adaptation of motor output, we use another metric called alignment index, defined as the temporal correlation between the measured hand forces and the commanded external force-field during the movement (see (Crevecoeur et al., 2020b, 2020a)). To focus on online adaptation, we calculated the alignment index over a time window from 100 ms after movement onset until movement end.

Previously (Crevecoeur et al., 2020b), we empirically showed that the temporal correlation between the hand force and external perturbation explained >90% of variance in hand acceleration, hence the unmodeled forces induced by the robot and sampling inaccuracies represented < 10% of unexplained variance. A high alignment index (value close to 1) is achieved when the temporal profile of the hand force is aligned with the external force-field disturbance. Importantly the measured force is the result of limb passive dynamics (arm mechanical impedance) and neural feedback, which cannot be dissociated. Thus, changes in temporal alignment can be interpreted as modulation in the feedback response under the assumption that the limb biomechanics was similar across conditions or experimental phases. This assumption is valid for the limb inertia as the configuration was the same across testing sessions. We show based on muscles activities that limb intrinsic stiffness and viscosity could be also considered constant as there was no systematic increase in baseline EMG activity, or coactivation of antagonist muscles (Figure 9).

#### Measure of anticipatory compensation (after effects)

To quantify anticipatory compensation, we measured the aftereffects in the null field trials that immediately follow a force field trial (see Figure 2B). In Experiment 1, we quantified after effect as the amount of lateral hand deviation in the opposite direction of the preceding force field. To ensure the measurements do not involve a significant online error compensation component, we quantified the hand deviation close to the start location at 1/3rd of the distance between the start and target locations, similar to the procedure described in (Crevecoeur et al., 2019).

In Experiment 2, we used error clamp trials (force channels) to restrict the hand movement to a straight line hence reducing the influence of online error compensation (Figure 2B). We then analyzed the lateral forces that participants generated in the force channel trials using standard procedure prescribed in (Smith et al., 2006; Coltman and Gribble, 2020). As a measure of the degree of anticipation in the force channel trials, we computed the anticipation index by estimating the slope of the relationship between the measured lateral force produced by the hand and the ideal force profile. A linear model with zero intercept was used to estimate the anticipation index on each force channel trial. The ideal force profile was calculated as the force profile that would have to be generated to fully compensate for the lateral forces had the rightward orthogonal force field been applied. If these forces were uncorrelated, the anticipation index was zero, and if the forces were identical the anticipation index was one.

### Statistical analysis

#### Experiment 1

In Experiment 1, the aim was to quantify the change in hand kinematics, forces and EMG activity participants across two conditions (HF and LF). Hence, to test the differences between within-participant means, we used paired two-sample t-tests. We did not need to correct for multiple comparisons in Experiment 1, as we only quantified the effect of the condition on the dependent variables, and thus only compared populations of values across conditions a single time (peak velocity, hand path error, aftereffects, peak force and alignment index). Statistical tests were considered significant at *p* < 0.05, although much lower p-values below this threshold were observed.

We measured the onset of changes in EMG responsible for changes in behavior across force field trials in the LF and HF conditions in Experiment 1. To contrast the differences between conditions, EMG data were averaged for each participant across the rightward orthogonal force field trials within LF and HF conditions. EMG averages were then collapsed into a 30-ms-wide (centered) sliding window, and paired t-tests were performed on the sliding window. We determined statistical significance as the first moment at which *p* < 0.005. It should be noted that corrections for multiple comparisons do not apply here for two reasons: the samples at each time step are involved in only one comparison, and consecutive samples are not statistically independent. Indeed, if there is a significant difference at a given time step, it is very likely that there is also a significant difference at the next time step because signals do not vary instantaneously. Hence, the risk of false-positive must not be controlled.

#### Experiment 2

In Experiment 2, we were interested in the effect of the trial number and condition (inconsistent vs consistent) on different variables of interest, that include peak velocity, hand path error, peak lateral force and force alignment index. We first confirmed the effect of task condition (inconsistent or consistent perturbation schedules) on the dependent variables by performing a paired two sample t-test between the total 40 force field condition in rightward direction in the inconsistent condition, and the last 40 rightward force field trials in the late consistent condition. Mainly, we fitted linear mixed models (LMMs) to determine the effect of trial number (‘*X_i,j_*’) on each dependent variable (*Y_i,j_*) of interest in the inconsistent condition. Here *i* represents the individual participant, *j* indexes the number of samples (trials) from a given individual. These models were fit using ‘*lme*’ library of ‘*Rstudio*’ using the following formula:

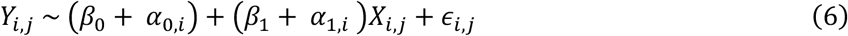

Intercept (*β*_0_) and the slope (*β*_1_) were fixed-effects reflecting the average population response that is constant across participant. Additionally, we included random effects that were different for each individual (*i*) to account for the idiosyncrasies. The random-effects include random offset (*α*_0*,i*_ ∈ *N*(0*, σ*_0_)) and random slope on the predictor (*α*_1_*_,i_* ∈ *N*(0*, σ*_1_)), where *σ*_0_ and *σ*_1_ are the variances of the random offset and slope variables and ∊*_i,j_* captures the residual. The linear mixed models were fitted with hand path error, peak force and alignment index as dependent variables, and we reported the F-statistics of the estimate as well as the corresponding p-value of the model. The effect of trial number was considered to be significant at *p* < 0.05 based on 1.96 ∗ *SD* from LMM.

We measured the onset of changes in EMG responsible for changes in behavior across early and late force field trials in the inconsistent condition in Experiment 2. Similar to the procedure used for comparing the EMG changes in Experiment 1, we used paired t-tests applied to a sliding time window on average participant data across orthogonal rightward force fields within early and late trials. We determined the strongest statistical significance across early and late trial EMG data as the first moment at which *p* < 0.005.

#### Multi linear regression analysis

To dissociate the contribution of different feedback control strategies to the observed hand path errors, we modeled lateral hand path errors across orthogonal force field trials in the unpredictable conditions of both experiments using a multiple linear regression with two predictors: peak lateral force and alignment index. The collinearity between the predictors has been quantified using Variation Inflation Factor (VIF). Model performance was assessed by computing the Pearson correlation coefficient between predicted and actual hand path errors across participants.

Additionally, we computed partial correlations using linear regressions with peak lateral force as the predictor while controlling for the alignment index, and evaluated the fit using the same procedure as for the full model.

To quantify the relative contributions of peak force and temporal force alignment, we extracted standardized regression coefficients from the multiple regression model. To further examine how these contributions evolved within a single force field trial, we fitted a sequence of multi-linear regression models predicting cumulative hand path error from the movement onset up to time *t*. For each model in the sequence, the predictors (peak lateral force and alignment index) were computed from the entire trial and remained constant, while the dependent variable – the cumulative hand path error – varied with time *t*. The resulting changes in standardized regression coefficients across time provide a measure of the relative importance of each predictor in explaining cumulative error within trial.

#### Unsupervised Clustering of participant force characteristics

We examined idiosyncrasies in how participants’ responses evolved across trials by applying an unsupervised clustering approach to classify participants into two groups. Specifically, we extracted coefficients ({β_0_, α_0_, β_1_, α_1_}) from the LMM, from Equation 6, where the peak force was modeled as a function of trial number in the inconsistent condition. These coefficients, representing both fixed effects (population-level intercepts and slopes (β_0_, β_1_)) and random effects (individual deviations (α_0,*i*_, α_1,*i*_)), were used as input features for K-means clustering. We visually noticed that the clustering algorithm divided participants mainly based on the slope of trial-by-trial change in the peak force.

This allowed us to classify participants into two groups based on how their peak force changed, effectively capturing both initial offset in responding to a novel force and how it evolved across trials. The clustering was performed in the multi-dimensional space defined by these LMM coefficients. To validate the clustering solution, we calculated the Silhouette score (Zaki and Wagner Meira, 2014), which quantifies how close a given participant to the others within the group subtracted from the average distance in the parameter space ({β_0_, α_0_, β_1_, α_1_}) from all the samples for another group. The Silhouette across participants is then averaged to get a group silhouette score. A positive group silhouette score indicates that participants were well grouped and meaningfully separated. The clustered groups had a Silhouette score of 0.46 indicating a meaningful separation.

After clustering participants based on their peak force responses in the inconsistent condition, we investigated potential differences between these two groups in other movement parameters across both inconsistent and consistent conditions, such as the modulation of hand path error and alignment index.

## Results

### Three different control strategies can influence motor responses to external perturbations

We begin by defining the three strategies previously identified in human reaching control (Figure 1). The first strategy is the classical trial-by-trial adaptation characterized by anticipatory production of a counter force (Figure 1B) to compensate for predictable disturbances when they are applied consistently across trials (Shadmehr, 2017). We will also refer to this strategy as “anticipatory compensation” (Figure 1B). The second strategy is robust control. It involves an increase in feedback gains to counteract unmodeled disturbances without prior knowledge of the perturbation profile (Gonzalez Castro et al., 2014; Franklin et al., 2017; Crevecoeur et al., 2019). This strategy results in faster movements and more vigorous responses to perturbations (Figure 1C). It is typically expected in inconsistent environments, where disturbances can be unpredictable. The third strategy is referred to as online adaptive control, and involves rapid adaptation of motor output during an ongoing movement. Behaviorally, this strategy can be characterized by a temporal re-alignment of the counter force produced in response to externally applied forces (Figure 1D). It is important to recall that the online adaptive control strategy is different from a standard feedback correction, as it involves a change in the shape of the correction, as previously reported in the context of force field adaptation (Crevecoeur et al., 2020c, 2020a). Observe that the last two strategies (robust and online adaptive control) are expected in inconsistent environments as they are in principle used when perturbations are not predictable, while anticipatory compensation is clearly expected in consistent environments.

Here, we document how the nervous system uses these strategies in concert to maintain successful reaching control across contexts and individuals. The remainder of the Results section is articulated as follows: we first concentrate on reaching control when perturbations are unpredictable. We show that these two strategies were evident in participants’ behaviour, that they are not trivially related to each other, and that their expression could be statistically associated with early or late phases of movement execution, respectively. Next, we measured the transition between inconsistent and consistent environments in Experiment 2 to highlight the contrast between the online mechanisms (robust and adaptive control), and the standard trial-by-trial anticipatory compensation strategy.

### Experiment 1: Concomitant Contribution of Robust and Adaptive Strategies in changing environment dynamics

We first sought to highlight robust and adaptive strategies in unpredictable environments. We contrasted the behavior of individual participants in two different environmental conditions (See Methods). These two conditions were labelled as low frequency (LF) and high frequency (HF) conditions based on how often perturbation trials occurred relative to the total number of trials (see Methods, Figure 2C).

In null field trials, participants reached the target relatively straight in both HF and LF conditions (Figure 3A). During unexpected orthogonal force field trials, the hand path deviated laterally in the direction of perturbation before the feedback correction could be observed (Figure 3A). At the group level, the hand deviated less in HF compared to LF condition during the force field trials (rightward and leftward) (Figure 3A right panel). Statistically, the hand path error, defined as the root mean square lateral deviation over the entire reach duration (see Methods), was significantly lower in HF compared to LF condition (paired t test: *t*_(16)_ = 5, *p* < 10^−4^).

This observed reduction in hand deviation could not be attributed to any anticipatory compensation since the perturbations were unpredictable in both conditions. We evaluated the after effect of a force field trial based on the hand deviation in the opposite direction during the subsequent null field trial close to the movement onset threshold (see Methods) (see Figure 3B). Across participants, this single exposure to a force field trial induced an after effect that was smaller in the HF condition (Figure 3B; paired t test: *t*_(16)_ = −3.2*, p* < 0.01). Smaller after-effects suggested that the trial-by-trial adaptation was less pronounced in the HF condition. Thus, when movement dynamics changed more frequently, participants were able to mitigate the impact of unpredictable disturbances by limiting the maximal and total travelled distances (Figure 3A), and reducing the after effect (Figure 3B) following the occurrence of a single force field trial.

Returning to the metrics of robust and adaptive control defined above, we found changes in hand velocity and force alignment that suggested a concomitant recruitment of both strategies. Firstly, the peak forward velocity was larger in the HF condition across the null field and force field trials, including after effect trials – defined as the first null field trial immediately following a force field trial (Figure 3C - E). Statistically significant differences in peak forward velocity between the HF and LF conditions were observed in force field trials (Figure 3C; paired t test: *t*_(16)_ = −5.2*, p* < 10^−4^), null (no perturbation) trials (Figure 3D; paired t test: *t*_(16)_ = −6.3*, p* < 10^−4^), and the after effect trials (Figure 3E; paired t test: *t*_(16)_ = −5.6*, p* < 10^−4^). Furthermore, participants applied larger peak lateral force in the HF condition to counteract both rightward and leftward force fields (figure not shown, *t*_(16)_ = 3.5*, p* < 0.005).

Overall, the general increase in hand velocity in the unperturbed direction across trials and larger forces in the perturbed direction were consistent with the upregulation of feedback control gains in the HF condition. Our interpretation is based on the fact that increase in forward velocity correlates with the vigor of feedback responses to perturbations, and both phenomena co-vary with the increase in uncertainty (Maurus et al., 2023), consistent with predictions of a robust feedback control strategy (Crevecoeur et al., 2019). Thus, the modulation of feedback gains is the hypothesis that best accounts for both behavioural measures. This strategy is unlikely to reflect predictive mechanisms because the perturbation induced by the force field increases with velocity, making it counterproductive. Notably, these results reproduce previous findings (Crevecoeur et al., 2019), now in a within-participants study, thereby showing how much participants could modulate control strength based on the frequency of perturbations.

Next, we assessed whether the participants also aligned their hand forces with the commanded force to better counteract the effects of the perturbations. The measured lateral force data from an exemplary participant illustrates the changes across HF and LF conditions qualitatively (Figure 4A, traces were normalized for illustration). In both HF and LF condition, the lateral measured hand force lagged behind the commanded force as a result of the unpredictability of force field. However, the shape of the measured hand force matched the commanded force better in the HF condition in both rightward and leftward force-field trials (Figure 4A). Such late temporal alignment of measured hand force has also been reported in previous independent studies (Crevecoeur et al., 2020a, 2020b). At the group level, we found that the correlation between the measured and commanded forces were significantly higher in the HF condition (paired t test: *t*_(16)_ = 4.1, *p* = 0.001). This stronger correlation demonstrates better temporal alignment of the feedback responses in the HF condition. Note that the temporal alignment index excluded the first 100ms of forces after movement onset (see Methods). Similar changes in temporal alignment between LF and HF condition were observed even when we computed correlation between forces restricted to the last 2/3^rd^ of each trial (paired t test: *t*_(16)_ = 3.5, *p* = 0.003).

**FIGURE 4.**
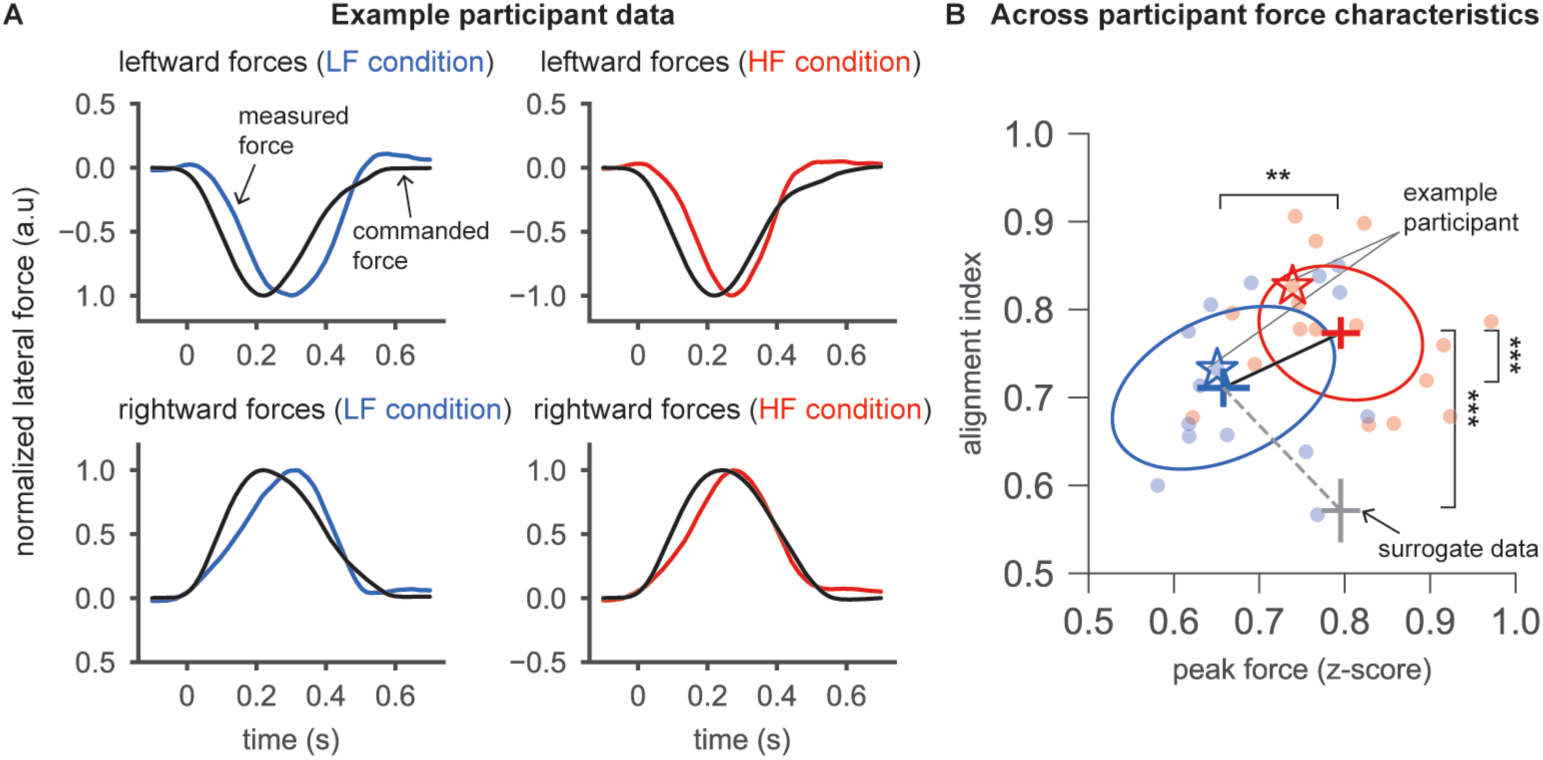
Concomitant increase in two feedback strategies from LF to HF condition. A) normalized lateral force profiles in rightward and leftward orthogonal force field trials of an exemplary participant, averaged across similar trial types within each of the HF and LF conditions. B) covariance ellipses depicting the variation in the alignment index and the peak force in HF and LF conditions. Each dot reflects one participant data in HF or LF condition. Star markers designate the exemplary participant from (A). Horizontal and vertical bars represent the S.E.M of peak force and alignment index respectively across participants (n=17). ***p<0.001, **p<0.01

Figure 4B illustrates that, in the HF condition, the participants concomitantly displayed significantly higher control strength as quantified by the peak force, and a higher temporal alignment of the force response with the external perturbation. To interpret these results in terms of distinct control strategies, it was necessary to verify that the increase in temporal alignment was not a trivial consequence of the increase in peak force. To rule out this potential confound, we generated a surrogate dataset where the measured lateral force produced by each participant (n = 17) in the LF condition was multiplied by a fixed gain such that the surrogate dataset had the same peak hand force as that of the corresponding participants in the HF condition. We observed that the temporal alignment of the surrogate dataset was lower than that of the participant data in both the LF (paired t test: *t*_(16)_ = 3.5*, p* < 0.005) and HF (paired t test: *t*_(16)_ = 9.5*, p* < 10^−6^) conditions (Figure 4B). This demonstrated that the alignment index in HF condition was not a trivial consequence of the increase in peak force, and it could therefore be attributed to a distinct modification of the reaching controller. This result corroborates previous simulation results (Crevecoeur et al., 2020b), showing that an increase in feedback gains did not lead to significant increase in the temporal alignment between measured and commanded forces. This argument was based on simulations in which a change in cost-functions aimed to mitigate the impact of the model errors did not increase the force alignment (see Figure 6C in (Crevecoeur et al., 2020b)). From a computational point of view, this difference indicates that evidence of changes in control strength – previously reported in (Gonzalez Castro et al., 2014; Crevecoeur et al., 2019) – possibly mediated by the recruitment of a robust policy or a change in the cost-function, can be dissociated from an adaptation of the internal model visible in the shape of the trajectory.

Thus, participants’ behaviour in Experiment 1 can be summarized as follows: there were concomitant increases in feedback gains (peak forward velocity and lateral force) and in the temporal alignment between measured and applied forces. These two aspects are compatible with robust and online adaptive feedback control, respectively. Since the same participants were tested in both contexts, we can deduce that each individual modulated their feedback control strategy in response to the likelihood of perturbations in the environment. Importantly, the reduction in after effect in the null trials that follow a force field trial suggested that the modulation of control responses could be dissociated from the classical trial-by-trial adaptation process. Our second experiment comes back on this distinction.

### Experiment 2: Dissociation between online strategies and trial-by-trial adaptation

In Experiment 2, we asked how the feedback responses evolved across trials when participants were exposed to a transition between unpredictable and predictable environments. Our goal was to highlight that the two feedback control strategies presented above could be behaviorally dissociated from the classical trial-by-trial adaptation based on anticipatory compensation.

Participants first performed reaching movements in a baseline condition in the absence of lateral perturbations. The baseline condition was followed by a highly inconsistent force field environment for several trials (referred as ‘inconsistent condition’), followed by a consistent environment where a single force field direction (rightward) persisted over many trials (see Methods, Figure 2B, D). We define consistency as the lag-1 autocorrelation between the external force-field parameters, describing how likely the force applied in a given trial persisted in the next one (Gonzalez Castro et al., 2014). To assess the amount of anticipatory compensation we used randomly interleaved error-clamp trials where force channels prevented lateral hand deviation, hence isolating the anticipatory responses from online feedback (Figure 2B). A scalar index of anticipatory compensation was computed by comparing the measured lateral hand force on the channel walls with the ideal force required if there was a force field in the rightward direction (see Methods). As expected, these measurements showed that the anticipatory compensation was absent in the inconsistent condition, then exhibited a gradual increase up to 80% in the consistent condition (Figure 5A). We verified that the average anticipation index across trials in the inconsistent condition was not significantly different compared to that of the baseline condition where no force fields were applied (paired t test: *t*_(16)_ = −1.4*, p* = 0.17).

**FIGURE 5.**
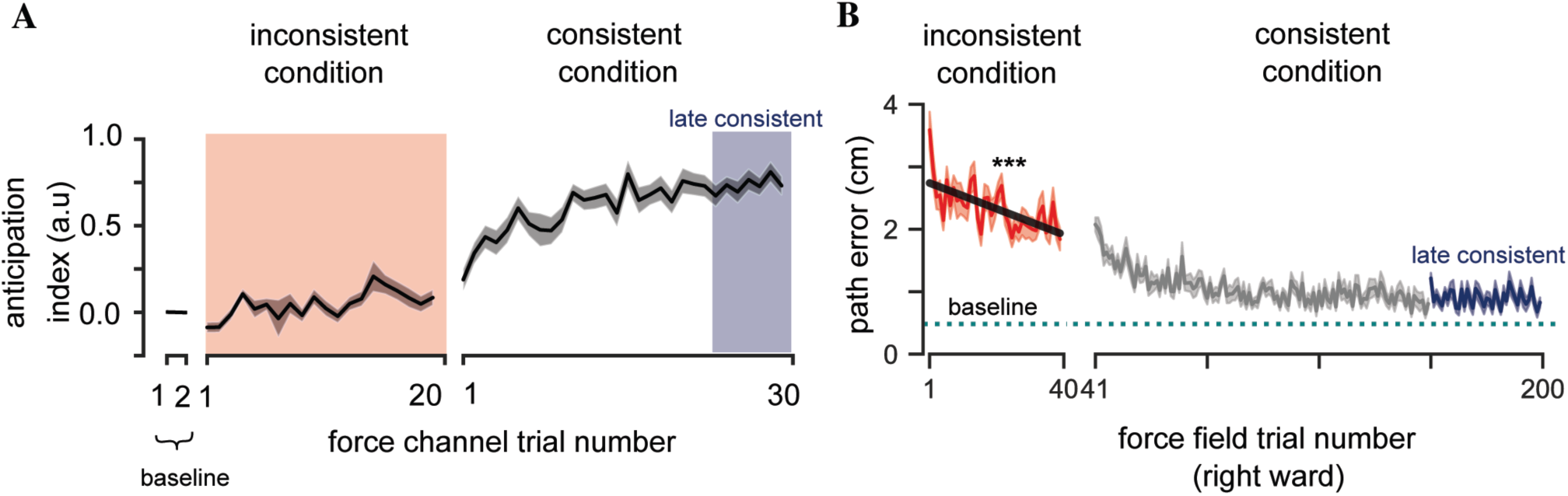
Decrease in movement error without anticipation. A) group average of anticipation index computed during force channel trials. B) group average of lateral hand path error from the rightward orthogonal force field trials across random and consistent force field condition. The average path error in the baseline trials is represented as a dotted line, and different task conditions are color coded. Asterisk symbols represent the statistical significance of the linear model fit to the trial by decrease in path error ***p < 0.001

Strikingly, the hand path error showed a strong and systematic trial-by-trial decrease in the inconsistent condition even though anticipatory compensation was clearly absent (Figure 5B). We fitted a linear mixed model on the hand path error as a function of the rightward force-field trial number (trial number from 1 to 40), while considering the participants as random effect (see Methods). The linear mixed model revealed a significant main effect of the trial number on the path error in the inconsistent condition (*F*_(1,636)_ = 33.2, *p* < 10^−5^). Once the environment switched to consistent rightward force field trials, the path error exhibited an exponential decay before stabilization (Figure 5B) close to 1 cm, which was slightly higher compared to baseline 0.5 cm (paired t test: *t*_(16)_ = 4.2*, p* < 0.001). Overall, to our surprise, approximately 65% of the overall reduction in path error occurred in the inconsistent condition alone, while the remaining 35% reduction occurred in the consistent condition. It is worthy to reiterate that participants’ behaviour in the inconsistent condition must have followed from adjustments in their online feedback responses, likely including a mix of robust and adaptive response as observed in Experiment 1. This interpretation is confirmed by our following analyses.

We examined the lateral hand forces following the same approach as in Experiment 1. Mainly, the temporal alignment between the measured lateral hand force and the commanded force improved during both inconsistent and consistent conditions, as shown in Figure 6A. This figure demonstrates that, across participants, the shape of the measured hand force matched better the commanded force in the first consistent trial compared to that of the first random trial (Figure 6A). Overall, at the group level, fitting a linear mixed model showed a significant effect of trial number on the temporal alignment in the inconsistent condition (*F*_(1,636)_ = 26.2, *p* < 10^−5^), as illustrated in Figure 6B.

**FIGURE 6.**
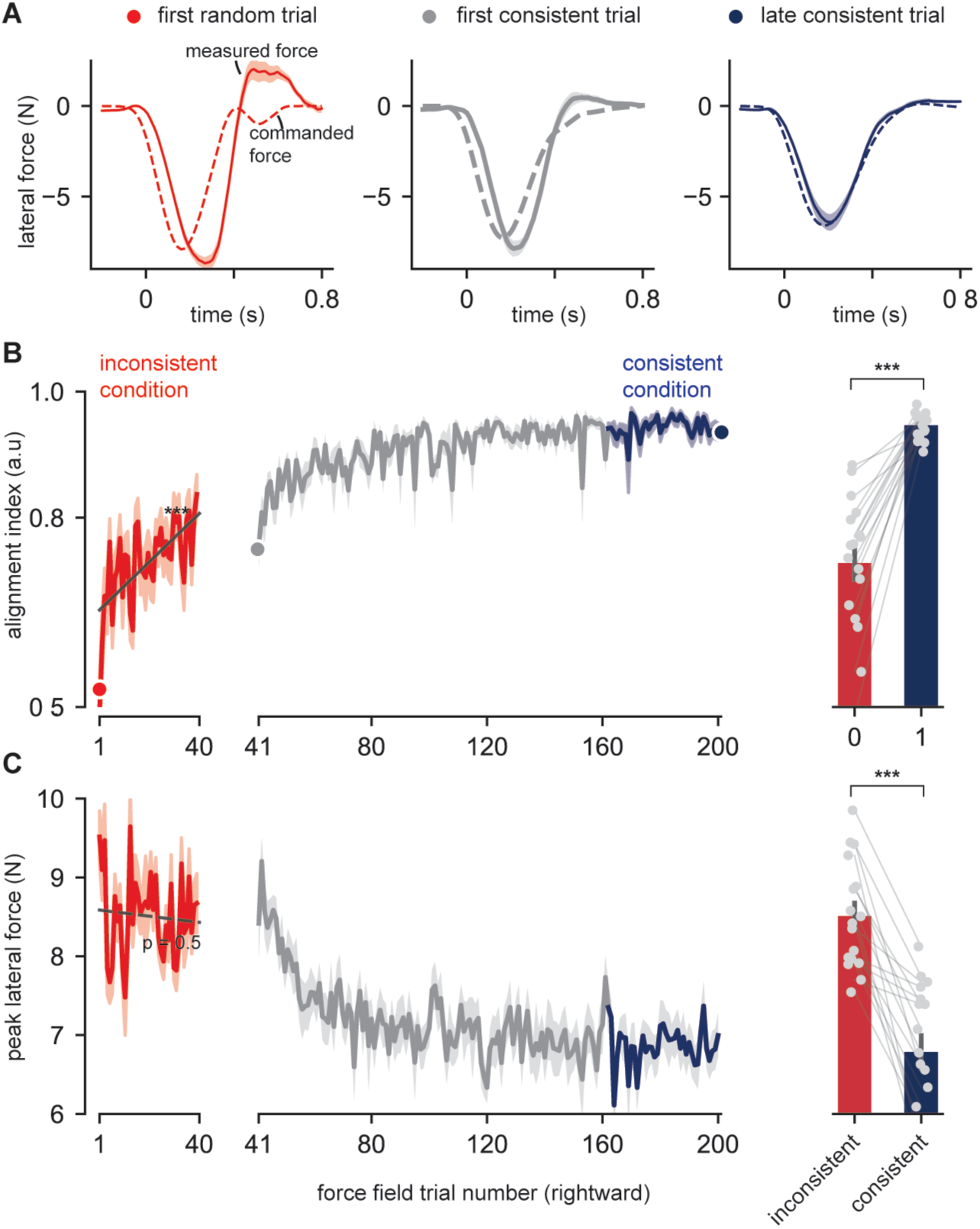
Trial-by-trial change in feedback responses. A) group level changes in lateral hand force for a given external force field disturbance in different trials B) group average of alignment index computed at each orthogonal rightward force field trial across random and consistent force field conditions. Bar plots depict the average across trials in random (red) vs late consistent (blue) conditions. C) group average of measured peak lateral force computed at each force field trial across inconsistent and consistent force field conditions. Bar plots depict the average across trials in random (red) vs late consistent (blue) conditions. Asterisks represent the statistical significance from paired t test with *** p < 0.001

By the last rightward force field trial in the consistent condition, this alignment was even larger, where the measured force was indistinguishable from the commanded force (Figure 6A). This high degree of alignment in the late consistent trials could be attributed to anticipatory force compensation, as anticipation reduces the mismatch between commanded and measured forces when the perturbations were predictable. On average, across participants and force-field trials, temporal alignment was higher in the late consistent condition (Figure 6B right panel; paired t test: *t*_(16)_ = −7.8*, p* < 10^−5^).

Besides the evolution of temporal alignment index, the measured peak lateral hand force was higher in the inconsistent condition, and remained high across trials until the beginning of the consistent condition (Figure 6C). Fitting a linear mixed model showed no effect of trial number in the inconsistent condition on the peak force (*F*_(1,636)_ = 0.3, *p* = 0.56). The peak lateral force (Figure 6C) in inconsistent condition was significantly higher compared to that of the consistent condition (paired t test: *t*_(16)_ = 7.3*, p* < 10^−5^), and similar patterns were observed in how peak forward velocity changed across force field and null field trials (figure not shown). This indicates that higher feedback gains were applied throughout the inconsistent condition, without a significant trial-by-trial change at the group level.

### Experiment 2: Quantification of individuals’ strategies

Here, we explored the possibility that individual participants exploited online control strategies to different extents. Almost all participants increased temporal alignment and decreased path error across trials. The key individual differences were in the modulation of the peak lateral forces. Remember that at the population level, the peak lateral force was on average constant across trials (Figure 6C; the line with almost zero slope, and *p* = 0.56). However, individual slopes and offsets from the linear mixed model fits varied widely across participants. The slopes of the peak lateral forces were either significantly positive or negative across participants (Figure S1A), while the slopes of the alignment index were positive, varying only in magnitude (Figure S1B).

A unimodal Gaussian mixture model fitted the distribution of participant-level slopes better than a bimodal model (BIC: −231 vs. −222.5; likelihood ratio test: LR = 8.14, *p* = 0.22), indicating that variation in control strategies was better described by continuum than qualitatively distinct subgroups (Figure S1C, D). Nevertheless, to characterize the two representative sides of this continuum, we used unsupervised clustering to classify participants into two groups based on their individual slope and offset (see Methods). This unsupervised clustering resulted in two groups with 9 and 8 participants. Notably, the clustering was similar even when using only the regression slopes because the two groups were separated by the sign of the slopes computed on the peak lateral forces.

Figure 7A-B illustrates how the peak lateral force and temporal alignment changed across trials in both groups. Similar to the pooled data from the entire population (presented in Figure 6B), the alignment between measured and commanded force increased across trials in both groups as well (Figure 7B; LMM on group 1: *F*_(1,348)_ = 30*, p* < 10^−5^; LMM on group 2: *F*_(1,304)_ = 76*, p* < 10^−5^), although participants in the group 1, referred to as “adaptive group”, displayed a larger alignment from the early trials.

**FIGURE 7.**
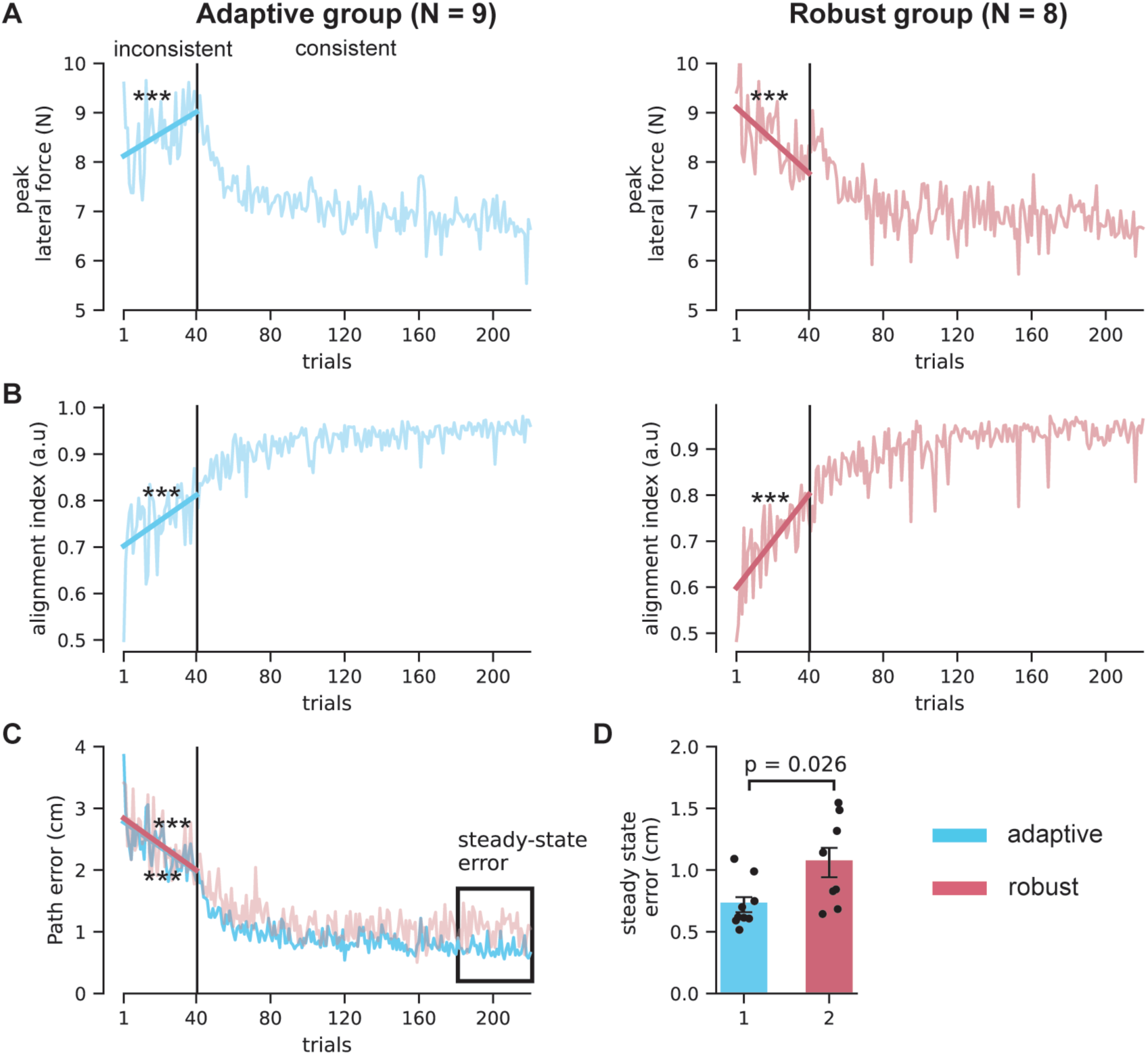
Individual differences in responding to inconsistent and consistent perturbations. A) change in peak lateral force across trials B) change in alignment index across trials. C) change in hand path error across trials. D) bar plots depicting the average steady state error of the last 40 trials between adaptive and robust participants groups, whereas the statistics represent independent samples t-test. Colored lines represent linear regressions of the dependent variable as a function of trial number. Statistical significance of the regression fit is depicted as asterisks, ***p<0.005, *p<0.05

Importantly, both groups exhibited stark differences in how the peak lateral force changed across trials in the inconsistent condition (Figure 7A). The adaptive group (Figure 7A, left, N=9) began with lower peak lateral forces that increased across trials (*F*_(1,348)_ = 17.8*, p* < 10^−5^;), and a comparatively high alignment index from the onset of the inconsistent condition. The second group (Figure 7A, right, N=8), referred to as the “robust group”, started with higher peak forces that decreased across trials (*F*_(1,304)_ = 31.6*, p* < 10^−5^), and lower initial alignment. These opposite trends in the evolution of the peak force across groups explain why the overall population level data showed no influence of trial number on this variable. Trial-by-trial changes in peak forward velocity closely paralleled the changes in lateral hand force.

Strikingly, the classification from the inconsistent condition predicted participants’ behaviour in the consistent condition. The adaptive group (high initial alignment index, trial-by-trial increase in peak forces) displayed lower steady-state movement errors after long exposure to consistent force field trials (Figure 7C, D). This was notable because participants were clustered into groups based on their feedback responses measured in the inconsistent condition and not based on the movement error data from consistent condition. Thus, there exists a relationship between the way they responded to unexpected disturbances and how they adapted to the force field in the consistent condition.

### Experiments 1 and 2: Influence of robust and adaptive feedback responses on motor errors

Here, we asked how robust and adaptive processes contributed to the changes in hand path in the unpredictable environments across the two experiments. We computed multi-linear regression models where the lateral hand path error of each participant, across orthogonal rightward force field trials in the unpredictable conditions, was fitted with a linear combination of two predictors: the measured peak lateral force and the temporal alignment index (Figure 8A, see Methods). The peak lateral force was used as a proxy of robust control, while the temporal alignment index provided a comprehensive metric for adaptive contributions. Although in general the force alignment depends on both anticipatory adjustments and online adaptation, its application to the unpredictable environments ensured that anticipatory adjustments remained negligible and therefore changes in this variable reflected online adaptation. We sought to examine the coefficients of this multi-linear regression to determine the influence of each predictor, namely the measured peak lateral force and the temporal alignment, on the hand path error as dependent variable.

**FIGURE 8.**
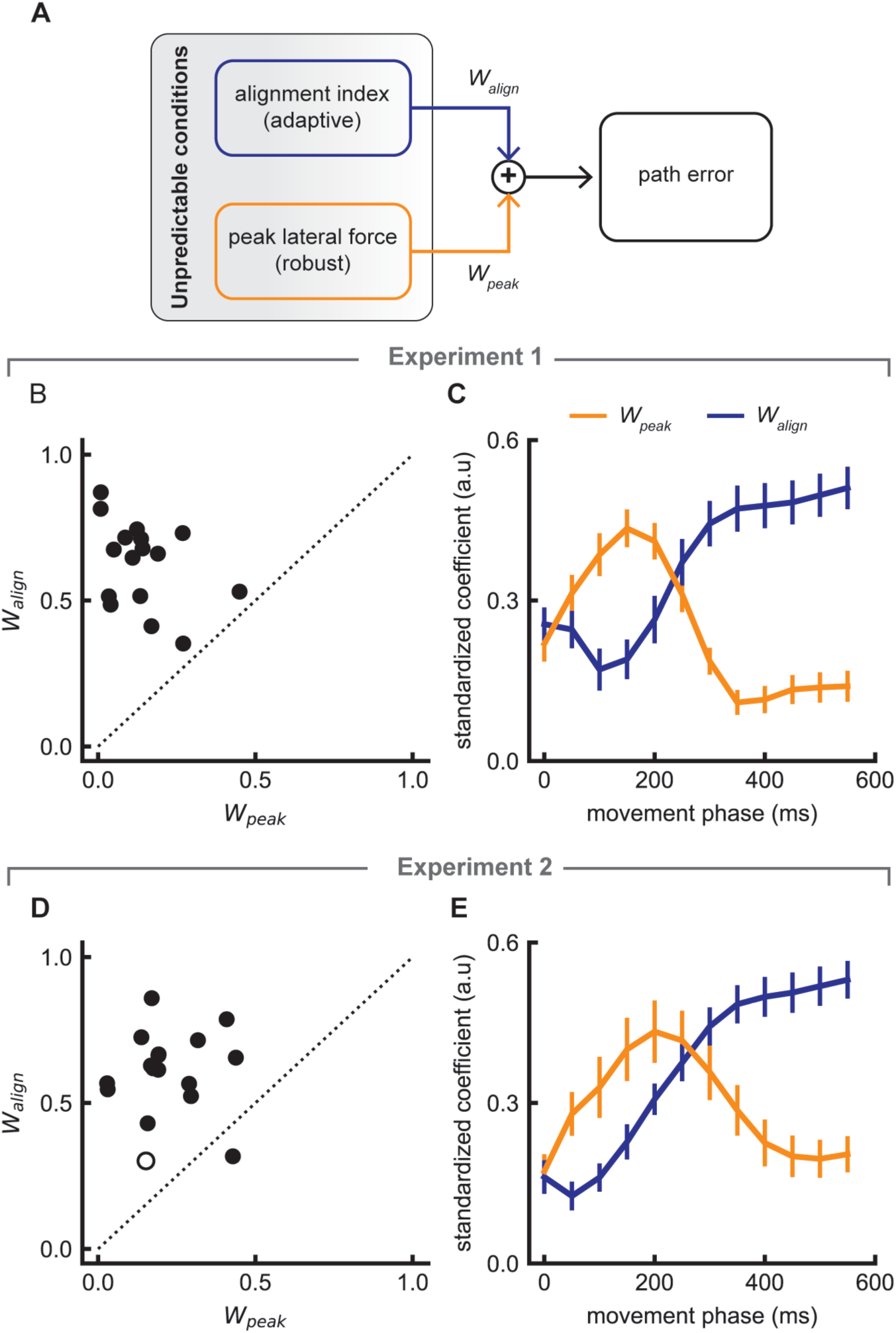
Contribution of different feedback strategies to the change in hand path error. A) Schematic to describe the multi-linear regression procedure B) Standardized coefficients (*W_align_* and *W_peak_*) of the corresponding predictor when the regressed variable was the path error in Experiment 1. C) Continuous change in the contribution of each predictor to the path error until the given time (movement phase) in Experiment 1. Note that the HF and LF conditions are merged here. D) same as (B) for Experiment 2. E) same as (C) for Experiment 2. Each dot represents the data from a single participant. Open circles represent participants for whom neither predictor yielded a statistically significant regression coefficient (p > 0.05). Errorbars represent ±*sem* around the mean across participants.

In Experiment 1, the relationship between the two predictors varied across participants. The cumulative distribution of Pearson correlation coefficients between peak lateral force and alignment index across participants is shown in Figure S2A. The average correlation coefficient (after Fisher z-transform) was r = 0.273 (SEM = 0.044), and the average variance inflation factor was 1.07, indicating low collinearity between the predictors. The multi-linear models of each participant predicted the path error well, with high correlation coefficients (average over individual participants after Fisher z-transformation: *r* = 0.66 in combined LF and HF conditions, p < 0.001). For each participant, partial correlation between peak force and path error, when controlling for temporal alignment, was much lower than the full model (average after Fisher z-transformation: *r* = 0.005, *p* = 0.92 compared to *r* = 0.66 for the full model; see Figure S2B), indicating a significantly higher effect of temporal alignment on the total hand path error. Figure 8B illustrates the standardized coefficients of regression from the multi-linear model, with data pooled from both HF and LF conditions. Each point corresponds to the regression coefficients of an individual participant. Clearly, across participants, the coefficients of the temporal alignment were larger, suggesting that the temporal alignment index had a larger impact on the hand path error.

We did a similar analysis on the data from the inconsistent condition of Experiment 2. The results of the multi-linear regression (Figure 8D) were qualitatively similar to those obtained in Experiment 1, even though the experiments were conducted on different groups of participants and with a different protocol. Like Experiment 1, the relationship between the two predictors varied across participants (Figure S2C). The average correlation coefficient (after Fischer z-transform) was r = 0.28 (SEM=0.042), with an average variation inflation factor of 1.08. For each individual participant the predicted output of the multi-linear regression models explained significant variations in hand path error (average after Fisher z-transformation: *r* = 0.66, p < 0.001). Partial correlation between peak force and path error, when controlling for temporal alignment, was lower than the full model correlation (average after Fisher z-transformation: *r* = 0.21, *p* < 0.001, compared to *r* = 0.66 for the full model; see Figure S2D), indicating that temporal alignment accounted for the majority of the relationship with path error. The regression coefficients corresponding to the temporal alignment were larger. Combined, the relative strength of coefficients from the multi-linear model and partial correlation analysis, in both Experiment 1 and 2, indicated that temporal alignment of lateral force had a larger influence on the hand path error than the peak lateral force.

To examine the possibility that robust and adaptive responses were recruited differently during movement, we further analyzed the standardized coefficients of each predictor by considering the cumulative path error until a given time horizon within the movement as a function of the peak force and temporal alignment indices (see Methods). We found that the change in peak force impacted the path error in the early movement phase until 250 *ms*, while the contribution of temporal alignment increased steadily throughout movement until the end, where it displayed a dominant effect (Figure 8C, E). Note that this result was remarkably consistent across Experiments 1 and 2.

Together, these results indicated that, in unpredictable environments, humans concomitantly exploited the modulation of control strength and the temporal alignment to mitigate the impact of unpredictable disturbances. Strikingly, each strategy influenced the hand path differentially depending on the phase of the movement: in the early movement phase (< 250 *ms* after movement onset), variations in peak force dominated the variation in hand path error, as the shape of the applied lateral force remained unchanged. As the movement progressed, the effect of the alignment index started to dominate after ∼250*ms* and became the main statistical predictor of the total path error.

### EMG responses emphasize the role of adjustments in feedback control

Finally, we examined whether participants adapted to disturbances by increasing limb mechanical impedance through co-contraction (Hogan, 1985; Burdet et al., 2001), or instead by adjusting online feedback responses. In Experiment 1, we analyzed differences in the shoulder muscle EMG activity (Pectorialis Major (PM) and Posterior Deltoid (PD)) during orthogonal force field trials between LF and HF conditions (Figure 9A, B). In Experiment 2, we compared the first four force-field trials with the last four force-field trials of the inconsistent condition (Figure 9C, D).

**FIGURE 9.**
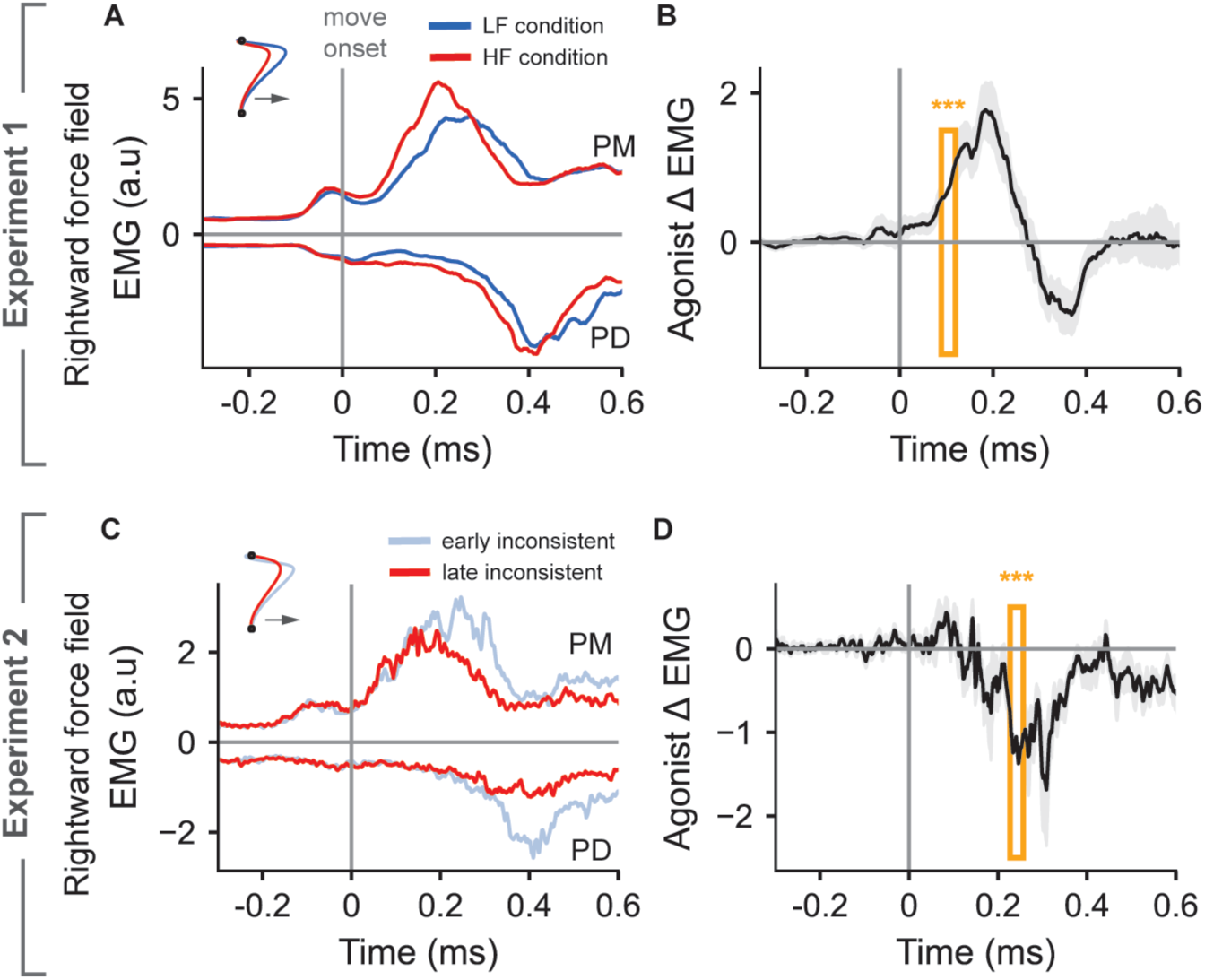
Surface EMG recordings from the Pectorialis Major (PM) and the Posterior Deltoid (PD) muscles that act as agonist and antagonist respectively for the rightward perturbation. A) group average PM and PD responses in HF and LF conditions in Experiment 1 B) the difference in the agonist (PM) activity between HF and LF conditions in Experiment 1. C) same as A) where cyan colored traces represent average of first four (early) trials and red traces represent average of last four (late) trials in the inconsistent condition in Experiment 2. D) same as (B) for the early and late trials in Experiment 2. Vertical gray line represents the movement onset time, computed as the time at which hand moves out of the start location. Shaded area represents the s.e.m around the mean value across participants. Three stars (***) represent the statistical significance from a running window paired t-test (see Methods) with **p < 0. 005**. Orange rectangle represents the first-time window at which the set statistical significance was attained.

In the rightward forcefield trials, no significant differences were observed in baseline co-activation of shoulder muscles prior to movement onset (−300*ms* to −100 *ms*) in either experiment. In Experiment 1, the baseline EMG activity did not differ between conditions for both muscles (PM: *t*_(16)_ = −0.2*, p* = 0.8; PD: *t*_(16)_ = 0.9*, p* = 0.3). Similarly, in Experiment 2, no significant baseline differences were found for the PM (*t*_(16)_ = −0.64*, p* = 0.5) or PD muscles (*t*_(16)_ = −0.94*, p* = 0.36). Consistent with these results, condition-dependent modulation was absent in leftward force field trials prior to movement onset in either experiment (Figure S3). These findings indicate that the observed modulation of reaching trajectories did not follow from any systematic increase in limb stiffness through co-activation of shoulder muscles prior to movement onset.

In contrast, both experiments showed a condition-dependent modulation of EMG activity during movement: In Experiment 1, the HF condition showed an increased agonist PM activation early in the movement, followed by reduction during the late movement period compared with the LF condition (Figure 9B). In Experiment 2, the late trials in the inconsistent condition showed a significant decrease in agonist PM activity during the late movement period (Figure 9D). Similar condition-dependent modulation of agonist PD activation during movement was observed in leftward force field trials across both experiments (Figure S3).

Additionally, comparing null-field (no perturbation) trials between the inconsistent and baseline conditions in Experiment 2 revealed no significant differences in the baseline EMG responses (−300 to −100 *ms* prior to movement onset) for either muscle (PM muscle: *t*_(16)_ = 0.9*, p* = 0.3; PD muscle: *t*_(16)_ = −1.9*, p* = 0.075) (figure not shown). These observations suggest that the behavioral modulation between baseline and inconsistent environments cannot be explained by systematic increase in co-activation in the PM and PD muscles, which could have impacted the mechanical stiffness of the shoulder joint.

## Discussion

This study highlights three components of human reaching control that are engaged to different extents dependent on the context. Besides standard trial-by-trial adaptation, we demonstrated that the CNS simultaneously employed two distinct feedback control strategies in unpredictable environments: the first is an increase in feedback gains, enhancing movement vigor and corrective forces, whereas the second is an online alignment of the temporal profile of the measured force with the force-field. These strategies were applied concurrently but produced dissociable effects on the movement trajectory, enabling performance improvements even when perturbations were unpredictable. Moreover, an individual’s preference in the unpredictable environment predicted their capacity to adapt in the consistent environment, offering new insights into the relationship between online control and trial-by-trial adaptation.

Motor adaptation has been typically studied using single metrics, such as maximum deviation (Thoroughman and Shadmehr, 2000; Scheidt et al., 2001), angular error (Morehead et al., 2017; Hadjiosif et al., 2021), or force in error-clamp trials (Smith et al., 2006). In contrast, we used multiple kinematic and force-based metrics to dissect a single reach into distinct control components. From recent work, we interpreted the peak velocity and corrective forces as proxies for robust control (Cluff et al., 2019; Crevecoeur et al., 2019; Maurus et al., 2023), whereas force alignment reflected online adaptive control (Crevecoeur et al., 2020b, 2020a). Lateral forces in channel trials served as a measure of anticipatory compensation following standard techniques. Previous simulations (Crevecoeur et al., 2019, 2020b) and our surrogate data showed that increasing feedback gains does not necessarily realign corrective forces. Thus, these two can be dissociated and attributed to distinct functions of the motor system. Both feedback gains and temporal alignment varied across contexts with different aftereffects, suggesting they are also dissociable from standard trial-by-trial adaptation measured in error-clamps. This approach captured the interplay between feedback-driven and anticipatory mechanisms by which the brain regulates movement trajectories.

Although we cannot definitively dissociate increased limb impedance from increased feedback gains, several observations support the latter. We observed no increase in co-coactivation of antagonist muscles in force-field environments compared to baseline, as expected if participants used impedance control (Hogan, 1985; Burdet et al., 2001; Calalo et al., 2023). In fact, average EMG activity was modulated during the movement. Stronger feedback corrections to lateral disturbances were paralleled by higher peak forward velocities in both experiments, indicating a general increase in control gains consistent with robust control rather a disturbance-rejection process based on limb intrinsic impedance.

Beyond robust control, which is a model-free strategy, our findings help reconcile conflicting views on the role of internal models in feedforward (anticipatory) and feedback control. Earlier studies suggested these processes share a common internal model, as changes in feedback corrections correlated with feedforward adaptation (Wagner and Smith, 2008; Ahmadi-Pajouh et al., 2012; Cluff and Scott, 2013; Albert and Shadmehr, 2016; Maeda et al., 2018; Coltman and Gribble, 2020). But, alternative evidence indicates they are dissociable in variable environments (Yousif and Diedrichsen, 2012; Gonzalez Castro et al., 2014; Kasuga et al., 2015). Our results show that force-field adaptation consists of two components: anticipation and alignment, both can improve temporal correlation between commanded and measured forces between trials (Joiner et al., 2017) or within a trial (Crevecoeur et al., 2020a; Mathew et al., 2020). When the force fields are consistent, online adaptation can serve as prior information to anticipate the force in the next movement. Conversely, when force-fields are inconsistent, feedback alignment can improve online independent of anticipatory compensation. Thus, internal models from online adaptation could be used in anticipatory control in context-dependent manner.

Our identification of distinct robust and adaptive phases within a single reach also offers a novel perspective on traditional two-component models. The classic Woodworth model (Woodworth, 1899; Elliott et al., 2001) posited a ballistic (open-loop) initial impulse followed by a corrective control phase. Similarly, Sainburg and colleagues (Sainburg et al., 1999) proposed a sequential link between a feedforward phase and a postural control phase during stabilization near the goal. However, our results suggest that, in a dynamic environment, early and late movement phases were governed by distinct feedback control strategies instead of discrete transitions from open-loop to closed-loop control. We propose that, in the presence of varying perturbations, the initial movement path reflected a robust strategy designed to handle unmodeled perturbations through high but costly control gains (Crevecoeur et al., 2019). As the movement progresses and sensory data accumulated, the system transitioned to online adaptation, exploiting updated internal models for more efficient corrections. Thus, our framework reinterprets the control of ongoing reaching as a transition between robust and adaptive feedback strategies, rather than a switch from open-loop to closed-loop control.

Our regression analyses confirmed that adaptive responses became the dominant predictor of hand path error at ≈ 250 *ms* after movement onset, consistent with previous estimates based on timing of EMG modulation (Crevecoeur et al., 2020a). Additional evidence from various tasks demonstrates that humans adjust control policies within a movement in response to changing dynamics and task demands (Braun et al., 2009; Kobak and Mehring, 2012; Crevecoeur et al., 2020a, 2020b; Comite et al., 2022, 2023; Kalidindi and Crevecoeur, 2023; Orban De Xivry and Hardwick, 2025). Collectively, these findings highlight the need to combine static models of sensorimotor control (Todorov and Jordan, 2002; Scott, 2004) with rapid and online update in model parameters and controller design (Kalidindi and Crevecoeur, 2023).

We highlighted individual differences in participants’ responses to unexpected perturbations that correlated with their ability to adapt in the consistent environment. Participants who responded to a novel force field by increasing feedback gains tended to exhibit initially lower force alignment, whereas those with lower feedback gains showed higher alignment at the beginning of the inconsistent environment (Figure 7). This dissociation allowed the unsupervised classifier to separate participants into two distinct groups: a “robust” and an “adaptive” group. Both groups achieved similar performances when repeatedly exposed to unpredictable perturbations, however they adjusted feedback gains and force alignment differently. A similar trade-off between robust and adaptive strategies has been reported previously (Cluff et al., 2019), and we believe that our observations hinged on the same underlying individual traits. Notably, the robust group also displayed lesser adaptation in the consistent scenario, leading us to hypothesize that these common aspects of control and adaptation may be directly linked to an individual’s ability to form an internal model of movement dynamics.

In light of the well-known trade-off between efficiency and robustness in control theory (Doyle, 1978; Zames, 1981), we speculate that the CNS regulates recruitment of these strategies based on the quality and adaptability of its internal models, combined with an evaluation of environmental uncertainty. When internal models are inaccurate and uncertainty is high, the CNS prioritizes stability by recruiting a robust strategy that mitigates worst-case performance despite higher corrective costs (Crevecoeur et al., 2019; Maurus et al., 2023). This could explain the behavior of participants from the “robust” group. Conversely, when internal models are more accurate and adaptable, the CNS may rely on rapidly adjusting internal representations with the risk of large initial errors. Across participants, this recruitment likely depends on a combination of personal traits and contextual factors that ultimately dictate how effectively a participant can reduce errors in a given environment.

Future research should investigate which neural circuits implement these strategies. Robust control through increased control gains has been associated with increased short- and long-latency feedback responses (Crevecoeur et al., 2019; Maurus et al., 2023, 2024; Dimitriou, 2018), suggesting modulation of spinal contributions through gain-scaling properties of the peripheral feedback loops (Pruszynski et al., 2009; Kurtzer, 2014; Guang et al., 2024). Adaptive control, in contrast, involves a state feedback controller (Izawa et al., 2008; Braun et al., 2009; Crevecoeur et al., 2020b), typically attributed to the long latency pathway – a distributed network encompassing primary sensory and motor cortices, premotor cortex, parietal regions, and the cerebellum (Kurtzer et al., 2008; Pruszynski et al., 2011; Omrani et al., 2016; Crevecoeur and Kurtzer, 2018; Kalidindi et al., 2021; Takei et al., 2021; Kalidindi and Crevecoeur, 2026). Online adaptation depends on continuous state-estimation error signal to update controller parameters (Crevecoeur et al., 2020b; Bicknell and Latham, 2025), critically requiring a common definition of state variables and error signals for both adaptation and control. Thus, regions capable of conveying error signals to the long-latency network – such as associative areas and the cerebellum – are prime candidates. Understanding how control and adaptation occur concurrently may help bridge theories of motor control (Guigon, 2023; Kalidindi and Crevecoeur, 2023) with motor adaptation (Shadmehr et al., 2010; Heald et al., 2021), which have largely been studied separately.

In conclusion, we propose a conceptual model in which the CNS combines a model-free robust strategy and a model-based adaptive strategy, with flexible recruitment depending on environmental statistics. The gradual transition between these strategies within a single movement clarifies how humans transition from model-free to model-based online corrections. By uncovering individual differences in adaptation to consistent environments, our results reveal a clear link between an individual’s feedback control responses and their capacity for motor adaptation across trials.

## Acknowledgements

HTK was supported by the Fonds de la Recherche Scientifique (FRS-FNRS) Chargé de recherche Grant CR 252 (FC 043127), and Research fellowship from Radboud Excellence Initiative. FC was supported by the FRS-FNRS Grant 1.C.033.18 (FC 036239).

## Supplementary Figures

**Figure S1.**
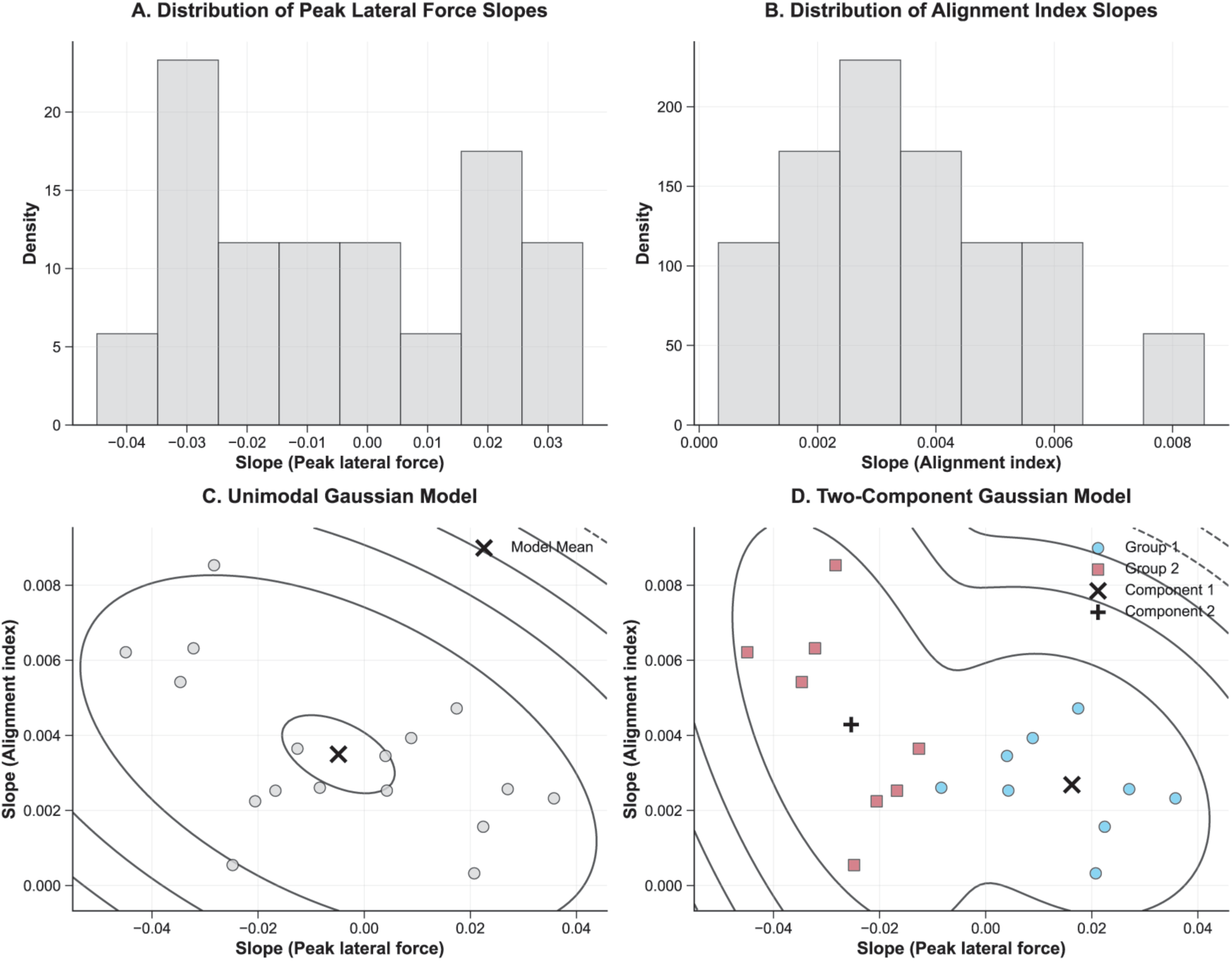
Distribution of slopes from linear mixed models. **A.** Slopes of peak lateral force across participants **B.** Slopes of alignment index across participants **C.** Unimodal gaussian mixture model fit on across participant slopes **D.** Bimodal (2-component) gaussian mixture model fit on the slope data. In the two-component model, the separation of groups in each mode (red and cyan colors) reflect the robust and adaptive groups found by the K-means clustering method in Figure 7.

**Figure S2.**
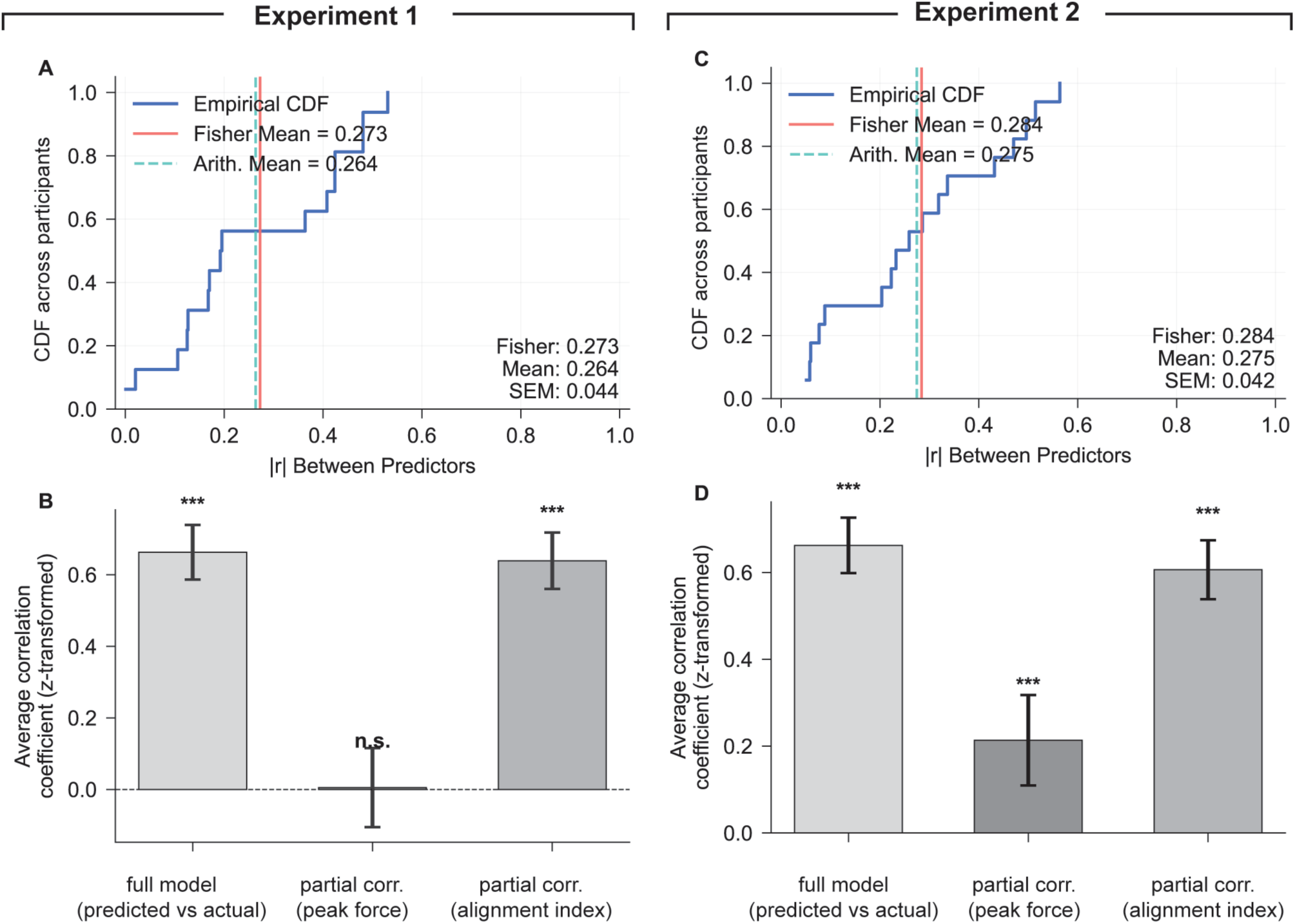
Multi regression analysis **A.** Cumulative density function of Pearson correlation coefficient between the predictors across participants in Experiment 1. **B.** Comparison between full regression model vs partial correlation analysis by controlling for each predictor in Experiment 1. **C.** Same as (A) for Experiment 2. **D.** Same as (B) for Experiment 2. Vertical lines on the top row subplots denote the mean of correlation coefficients across participants. ***p < 0.005, ^n.s^ p > 0.05

**Figure S3.**
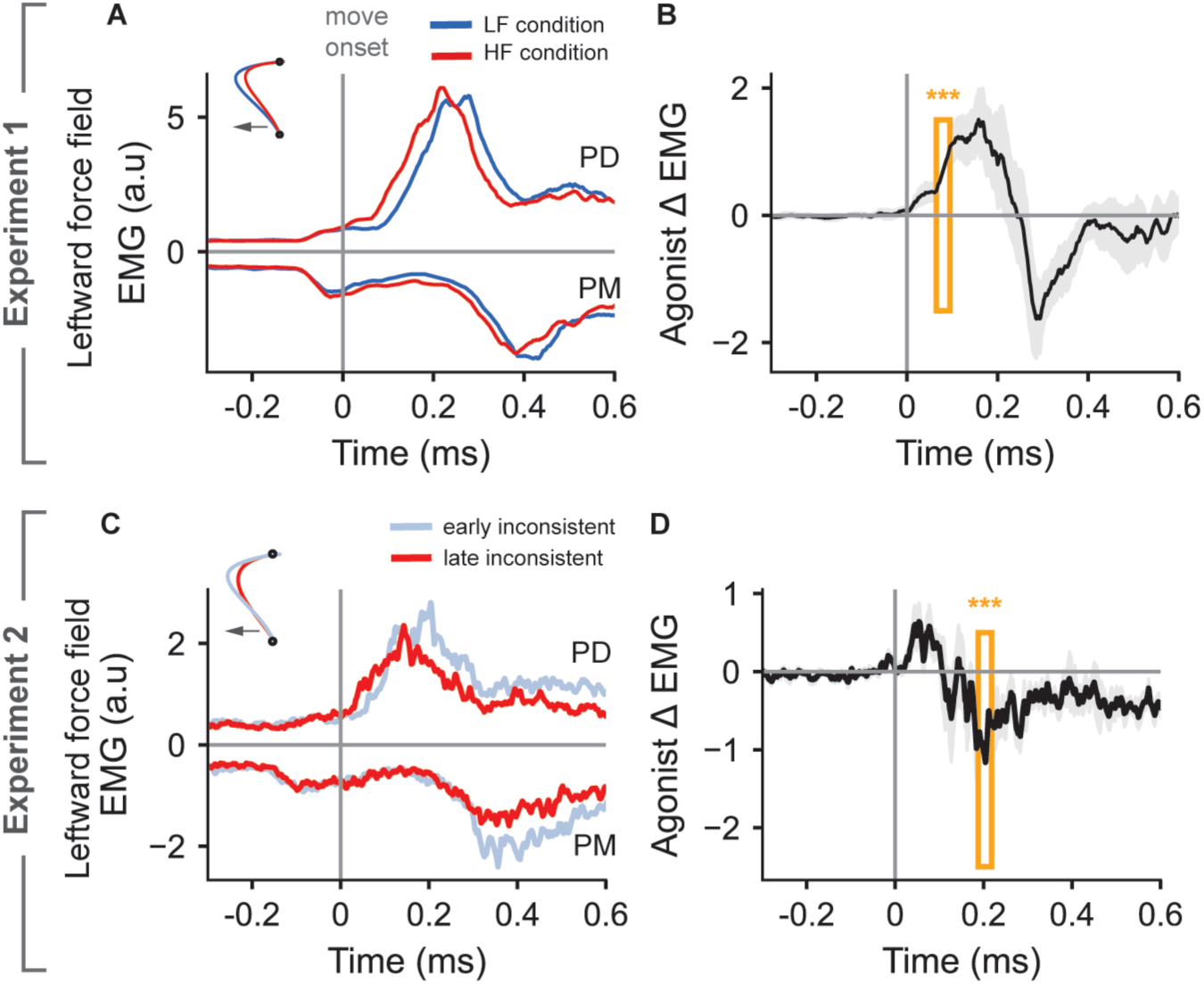
Surface EMG recordings from the Posterior Deltoid (PD) and the Pectorialis Major (PM) muscles that act as agonist and antagonist respectively for the “leftward orthogonal force field perturbation”. A) group average PD and PM responses in HF and LF conditions in Experiment 1 B) the difference in the agonist (PD) activity between HF and LF conditions in Experiment 1. C) same as A) where cyan colored traces represent average of first four (early) trials and red traces represent average of last four (late) trials in the inconsistent condition in Experiment 2. D) same as (B) for the early and late trials in Experiment 2. Vertical gray line represents the movement onset time, computed as the time at which hand moves out of the start location. Shaded area represents the s.e.m around the mean value across participants. Three stars (***) represent the statistical significance from a running window paired t-test (see Methods) with p < 0.005. Orange rectangle represents the first-time window at which the set statistical significance was attained.

