## Supplementary Material for "When anticipation is not enough: a mixture of robust and adaptive feedback control strategies improve reaching in dynamic environments"

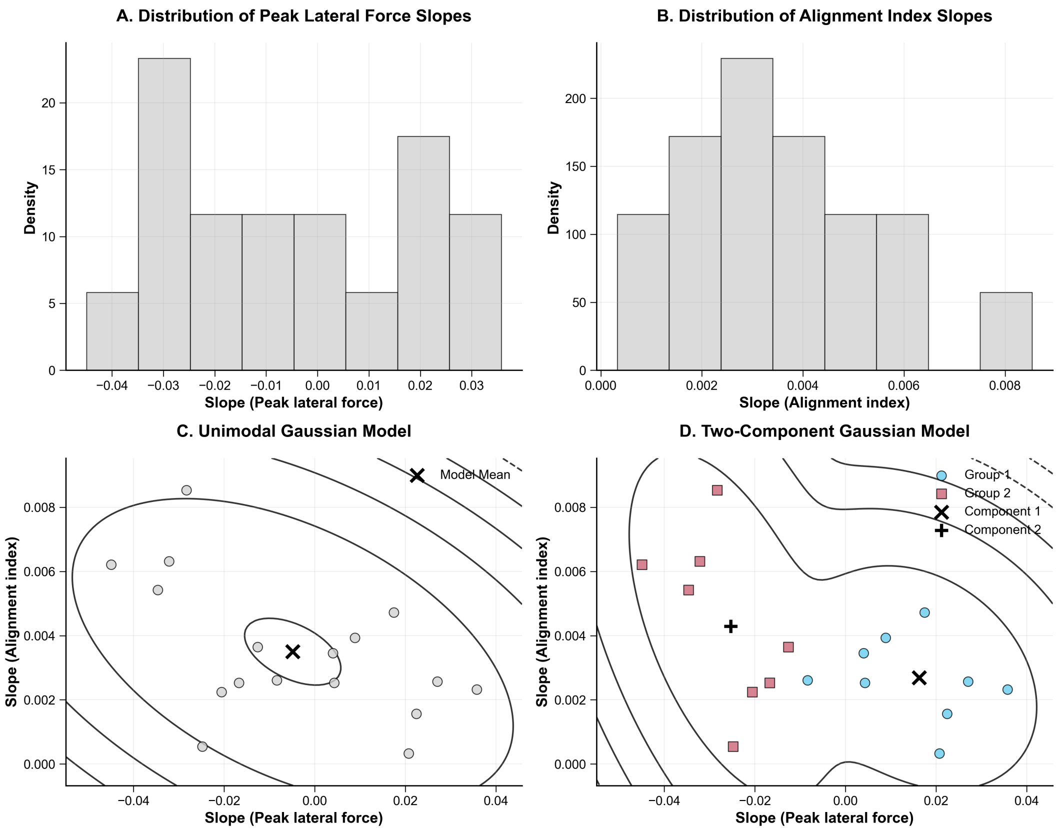


**Figure S1.** Distribution of slopes from linear mixed models. **A.** Slopes of peak lateral force across participants **B.** Slopes of alignment index across participants **C.** Unimodal gaussian mixture model fit on across participant slopes **D.** Bimodal (2-component) gaussian mixture model fit on the slope data. In the two-component model, the separation of groups in each mode (red and cyan colors) reflect the robust and adaptive groups found by the K-means clustering method in Figure 7.

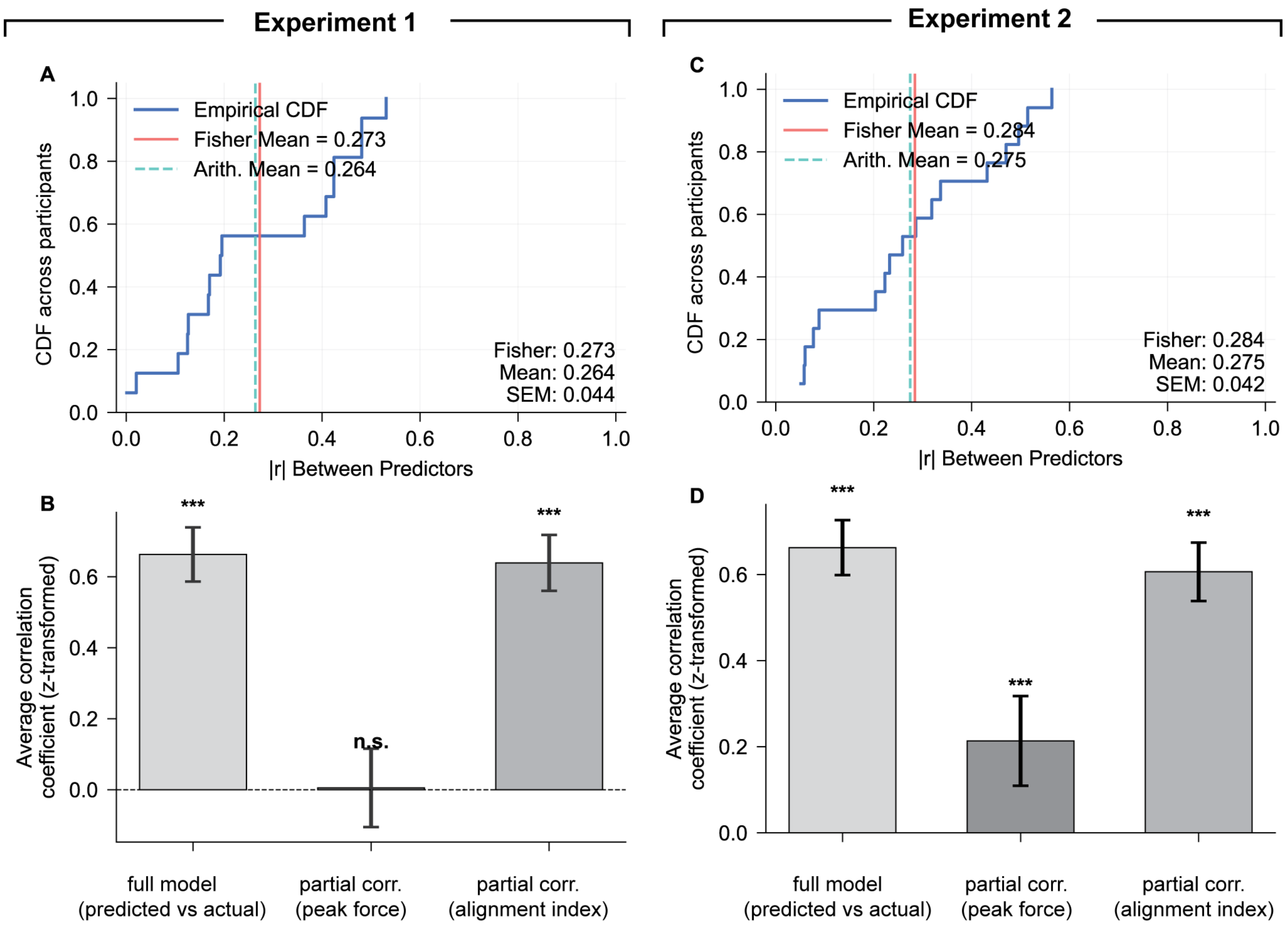


**Figure S2.** Multi regression analysis **A.** Cumulative density function of Pearson correlation coefficient between the predictors across participants in Experiment 1. **B.** Comparison between full regression model vs partial correlation analysis by controlling for each predictor in Experiment 1. **C.** Same as (A) for Experiment 2. **D.** Same as (B) for Experiment 2. Vertical lines on the top row subplots denote the mean of correlation coefficients across participants. ***$p < 0.005$, $^{n.s}$ $p > 0.05$

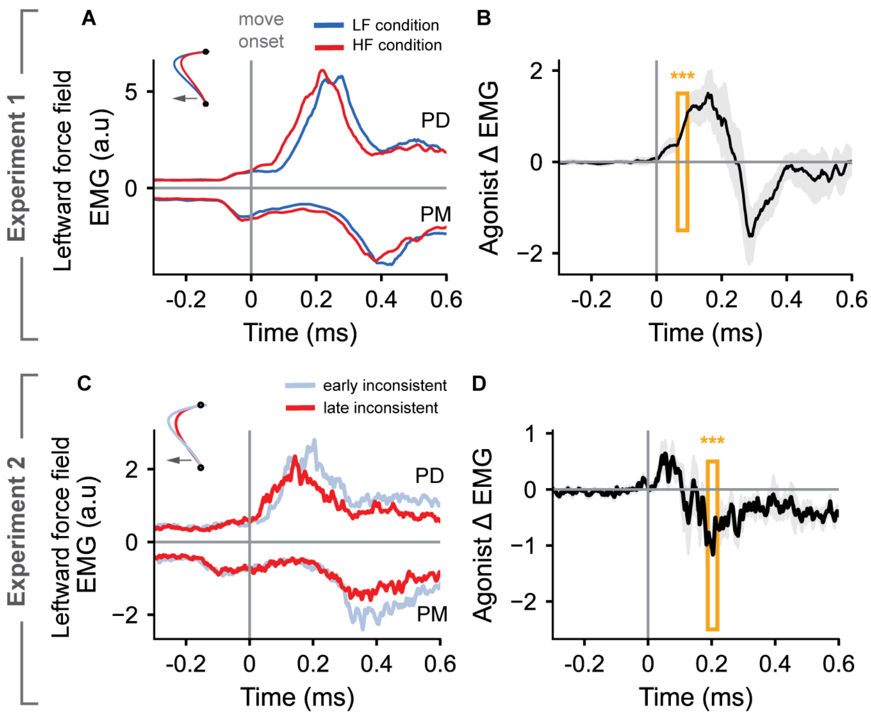


**Figure S3.** Surface EMG recordings from the Posterior Deltoid (PD) and the Pectorialis Major (PM) muscles that act as agonist and antagonist respectively for the "leftward orthogonal force field perturbation". A) group average PD and PM responses in HF and LF conditions in Experiment 1 B) the difference in the agonist (PD) activity between HF and LF conditions in Experiment 1. C) same as A) where cyan colored traces represent average of first four (early) trials and red traces represent average of last four (late) trials in the inconsistent condition in Experiment 2. D) same as (B) for the early and late trials in Experiment 2. Vertical gray line represents the movement onset time, computed as the time at which hand moves out of the start location. Shaded area represents the s.e.m around the mean value across participants. Three stars (***) represent the statistical significance from a running window paired t-test (see Methods) with $p < 0.005$. Orange rectangle represents the first-time window at which the set statistical significance was attained
